# Plant pathogenic *Ralstonia* share two core methyl-accepting chemoreceptors that drive chemotaxis toward distinct amino acid profiles

**DOI:** 10.1101/2025.04.07.647700

**Authors:** JK Avalos, VN Elmgreen, CR Guerrero, KJ Grulla, K Miqueo, RE Parales, TM Lowe-Power, RA Schomer

## Abstract

*Ralstonia solanacearum* species complex pathogens cause bacterial wilt disease in diverse plant families. These pathogens use chemotaxis and motility to discover host roots. The specificity of this process is conferred by methyl-accepting chemotaxis proteins (MCPs). We explored pangenomic variation of MCPs and other chemotaxis machinery across *Ralstonia* wilt pathogens. We classified 19 MCPs as broadly conserved, core MCPs, and we identified several accessory MCPs. Within their periplasmic sensing domains, two of the core MCPs contain a motif that is known to bind amino acid ligands: McpA1 and McpA2. To identify the ligands of McpA1 and McpA2, we constructed HyChemosensor strains. In HyChemosensor strains, the periplasmic sensing domains of the MCPs were translationally fused to the signaling domain of a model two component sensor, NarQ. The HyChemosensor assays revealed that the receptors, McpA1 and McpA2, have overlapping binding profiles for acidic amino acids. Nevertheless, McpA2 recognized a broader profile of amino acids. Quantitative swim plate assays confirmed that these receptors facilitate *Ralstonia*’s chemoattraction to amino acids. Consistent with prior evidence in a different *Ralstonia* strain background, *mcpA1* was dispensable for virulence on tomato. However, single or double mutants lacking *mcpA2* demonstrated reduced virulence on tomato following naturalistic soil soak inoculations. Thus, McpA1 and McpA2 have distinct roles in plant colonization, despite redundancy in their chemical specificity.

## Importanc

Chemotaxis is a widely conserved behavior that influences the ability of bacteria to navigate their environments and colonize hosts. Redundant and overlapping chemical specificities of MCPs complicate our ability to identify critical chemical signals and receptors in host-microbe interactions mediated by chemotaxis. We identified two chemoreceptors that have overlapping chemical-binding profiles but distinct roles in host colonization. These results emphasize the complexity of chemotaxis behavior in *Ralstonia* wilt pathogens. Our pangenomic analysis and HyChemosensor assays set the stage for deciphering how MCP-mediated signaling contributes to host colonization.

## Introduction

The *Ralstonia solanacearum* species complex are soil-borne bacterial wilt pathogens that cause disease in a wide range of botanical families (1). This species complex is genetically diverse and is composed of three species: *R. solanacearum* (phylotype II)*, R. pseudosolanacearum* (phylotypes I and III), and *R. syzygii* (phylotype IV), hereafter collectively called “*Ralstonia”. Ralstonia* survive in soil, where they can invade roots and spread systemically in the water-transporting xylem vessels of the plant, ultimately leading to fatal wilt disease. To complete their life cycle, *Ralstonia* must locate the host root, attach to the host root, and invade the root. Root exudate chemicals are a critical mechanism of dialogue between plants and soil microbes. These exudates alter soil pH to solubilize nutrients, deter growth of competing plants, and recruit microbes to the rhizosphere (2). *Ralstonia* uses chemotaxis to locate host roots (3, 4). Chemotactic behavior allows a bacterium to detect a chemical signal such as a root exudate compound and swim in a biased way, guided by the concentration gradient of that chemical.

Bacterial chemotaxis requires a conserved set of cytoplasmic proteins (CheA, CheB, CheR, CheW and CheY) and a suite of transmembrane chemical-sensing proteins called methyl-accepting chemotaxis proteins (MCPs). MCPs contain at least two functional protein domains: (1) a highly divergent, periplasmic sensing domain that binds ligands (e.g., 4-helix bundle, double cache, single cache, etc.) and (2) a conserved, cytoplasmic signal transduction domain. Many MCPs also contain a structurally conserved transmembrane HAMP domain, named for its presence in <u>H</u>istidine kinases, <u>A</u>denyl cyclases, <u>M</u>ethyl-accepting proteins and <u>P</u>hosphatases. MCPs directly detect the presence of different chemicals via the periplasmic ligand binding region (5, 6). Binding of ligands to the MCP periplasmic domain triggers the cytoplasmic methyl-accepting signal transduction domain to activate autophosphorylation of CheA (6, 7). CheA/CheY is cytoplasmic two-component phosphorelay system (8). Ultimately, CheY directly interacts with the flagellar motor to control the direction of flagellar rotation. Depending on the direction of flagellar rotation, the bacterium swims smoothly in one direction or the bacterium stalls in a “tumble”. The net result is that bacteria collectively swim up chemical gradients of attractants or down a gradient of repellents. CheB and CheR are involved in the methylation/demethylation of the MCPs, which tunes the signal, allowing the bacterium to adapt to changing concentrations of attractants (9, 10). Che signaling components are typically conserved across chemotactic bacteria, but the number of MCP chemosensors and the repertoires of chemicals that they detect varies widely among bacterial strains (11, 12).

*Ralstonia* mutants lacking cytoplasmic chemotaxis machinery (Δ*cheW* or Δ*cheA*) or flagella (Δ*fliC*) have virulence defects when challenged to locate tomato roots by naturalistic soil soak inoculations (4, 13). In contrast, the same mutants are fully virulent when allowed to bypass the roots via a stem inoculation (4, 13). Soil-dwelling and plant-associated bacteria including *Ralstonia* often encode dozens of MCPs in order to sense and respond to highly complex environments (11, 12, 14, 15). Only a few of the *Ralstonia* MCPs have been studied. To date, only an L-malate-binding MCP (McpM) is known to quantitatively contribute to virulence on tomato plants (16). MCPs that bind citrate (McpC and McpP), L-tartrate (McpT), borate (McpB), and amino acids (McpA) have been characterized, but mutants with individual knockouts of these MCPs were fully virulent (16–18). Like other tactic bacteria, *Ralstonia* strains can also sense chemical attractants indirectly through “energy taxis” (also called “aerotaxis”) (3, 19, 20). In energy taxis, bacteria sense optimal metabolic conditions through MCP-like proteins that have Per-Arnt-Sim (PAS) domains. PAS domains measure the physiological state of the cell rather than binding to a specific chemical ligand (21). *Ralstonia* strains have two well-characterized energy taxis proteins (Aer1 and Aer2) that additively contribute to pathogenic fitness when cells are challenged to locate tomato plant roots (3). Functional redundancy of MCPs is common in soil bacteria like the pseudomonads (22, 23). In some cases, multiple MCP gene deletions must be generated to determine whether an individual receptor detects a certain ligand (22–24). Alternate approaches that bypass the need for multiple mutations include isothermal titration calorimetry (ITC) on purified and often truncated proteins, or the generation of biosensor strains with hybridized chemoreceptors and two-component systems (25).

Although Aer1, Aer2, and McpM quantitatively contribute to virulence, their additive contributions do not fully account for the reduced virulence observed in non-tactic mutants (3, 16). Despite the increasing availability of *Ralstonia* genomic data, most MCPs remain uncharacterized. In this study, we used bioinformatics to infer evolutionary patterns of *Ralstonia* MCPs and identified MCPs that contained a known amino acid-binding motif: the previously characterized McpA (now “McpA1”) and the uncharacterized McpA2. We investigated the ligand specificity and biological role of these receptors in chemotaxis and in infection of tomato plants, revealing that amino acids function as chemoattractants that enable *Ralstonia* to locate host plants.

## Results

### The *Ralstonia* pan-genome contains 19 core chemoreceptors and at least two lineage-specific, accessory chemoreceptors

Previously, 22 putative MCP paralogs have been identified in the genome of the model strain *R. pseudosolanacearum* GMI1000 (phylotype I) and were confirmed to have orthologs in the related phylotype I strain Ps29 (16). To explore the genetic conservation and natural variation of *Ralstonia* chemosensing, we investigated the presence/absence of the 21 MCP paralogs that are associated with flagellar chemotaxis (16) (Fig. 1 & S1, Table S1), excluding the PilJ chemoreceptor that is associated with pilus-mediated twitching motility. We used BLASTP searches to detect these genes across a set of 106 diverse, high-quality *Ralstonia* genomes. This analysis revealed that 19 of the *mcp* genes are broadly conserved across the *Ralstonia* pangenome, which we hereafter call the “core MCPs”. We infer that all 19 core MCPs are vertically inherited because synteny analyses show that these genes are located in conserved neighborhoods (Fig. S2). Additionally, we identified two accessory *mcp* genes (RSp1363 and RSc1950) that are present in some of the lineages. Accessory *mcp* RSp1363 was present in phylotype I and a major branch of phylotype IV (IV-sequevar 8), and accessory *mcp* RSc1950 was present and predicted to encode a full-length 1,011 bp MCP in phylotype IIA (strains) and as a putative pseudogene of 990 bp in all phylotype I strains.

**Figure 1.**
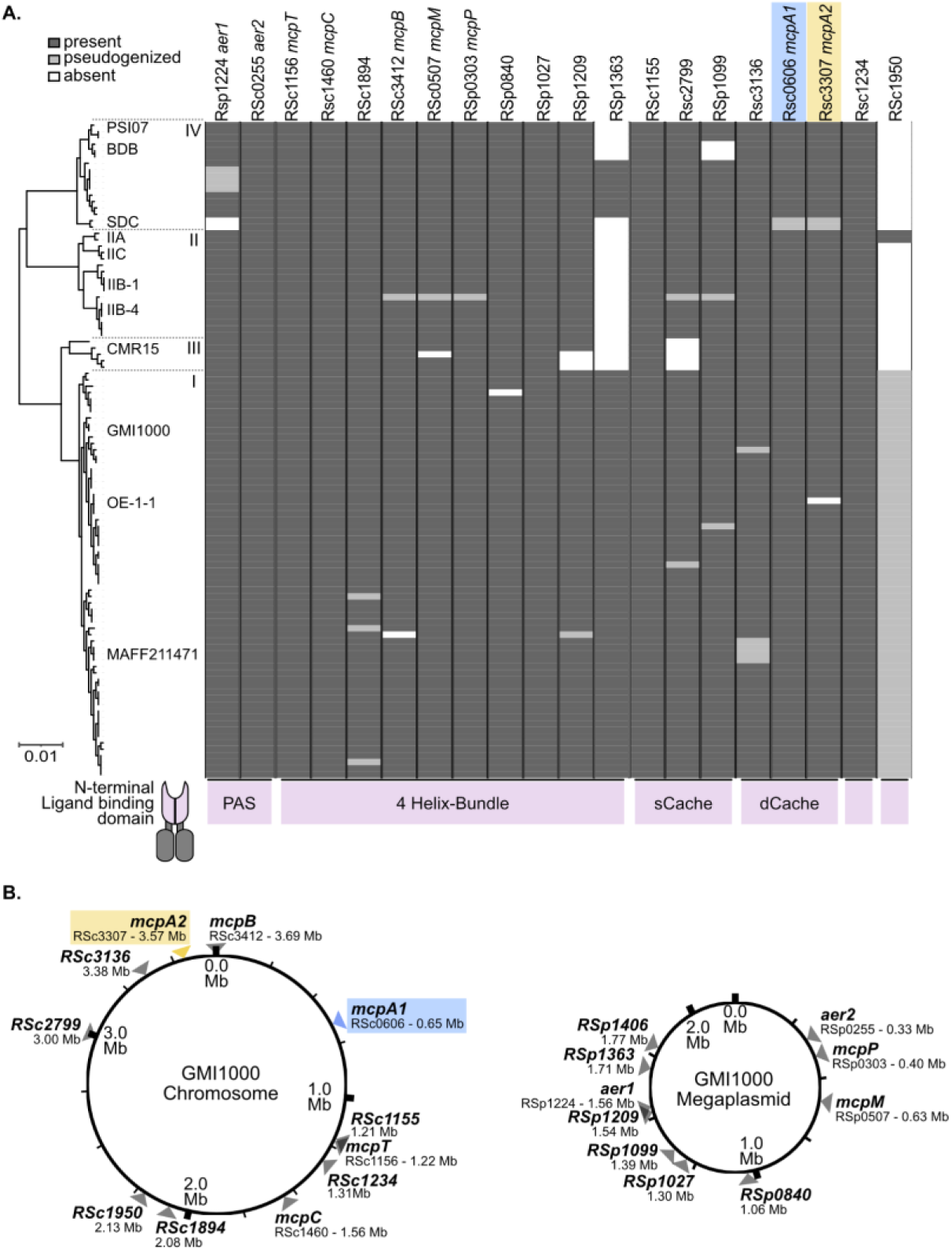
*Ralstonia* wilt pathogens have a large set of core MCP chemosensors and occasional accessory MCP chemosensors. (A) Presence/absence of orthologous MCPs across diverse *Ralstonia* genomes. The tree was built with the KBase SpeciesTree app, which creates a Multiple Sequence Alignment of 49 conserved bacterial genes and builds the tree with FastTree2. Phylotype I-IV are labeled on the tree. Presence/absence of *mcp* genes was determined through a combination of BlastP and synteny analyses. The type of N-terminal, periplasmic ligand binding/ sensing domain is indicated below each *mcp* paralog. Unlabeled domain indicates unclassified ligand binding domain types . (B) *mcp* genes are distributed across the chromosome and megaplasmid. Locations of each *mcp* in the genome of model strain GMI1000 are shown. Figure S1 shows the annotated tree with each genome labeled.

To sense diverse environmental stimuli, bacterial MCPs are modular and have variable N-terminal periplasmic sensing domains. We investigated the domain architecture of the core *Ralstonia* MCPs (Fig. 1A). The most common domain type is the 4-helix bundle (4HB) domain (PFAM12729), which is present in 11 of the MCPs. Six MCPs contained either single cache (PFAM17200) or double cache (PFAM08269) domains. The 2 MCPs previously identified as energy taxis sensors contained PAS domains (PFAM13426)(3). Interestingly, 2 MCPs had novel N-terminal domains that were not classified by InterProScan.

### Genes encoding MCPs are distributed across both the chromosome and megaplasmid while genes encoding other flagella and chemotaxis functions are enriched on the megaplasmid

To investigate whether the *mcp* paralogs are encoded in genomic hotspots, we investigated their locations in the bipartite genome of model strain *R. pseudosolanacearum* GMI1000. In most cases, the *mcp* genes locations are dispersed, but two *mcp* genes (RSc1155 and *mcpT*/RSc1156) are encoded sequentially in a putative operon on the chromosome. Out of the 21 *mcp* genes, 11 are encoded on the chromosome and 10 are encoded on the megaplasmid (Fig. 1B).

In contrast, most other flagellar and chemotaxis machinery is encoded on the megaplasmid. Genes involved in flagellar synthesis (*flgNMABCDEFGHIJKL* and *fliRQPONMLEFGHIJK*) and core chemotaxis machinery (*cheAWZBR* and two *cheY*-like genes) are located on the megaplasmid as gene clusters with conserved organization (Fig. S3-S5). Although most flagellar and chemotaxis genes are present in single copy, *Ralstonia* genomes encode multiple CheY and CheZ paralogs. Two *cheY*-like genes (RSp1401 and RSp1409) and one *cheZ*-like gene (RSp1402) are encoded in the *cheA*-containing chemotaxis cluster. Additionally, *R. solanacearum* and *R. syzygii* genomes contain another *cheYZ* locus on the megaplasmid (RALBFv3_RS23105-RALBFv3_RS23110) while *R. pseudosolanacearum* genomes contain another *cheYZ* cluster on the chromosome (RSc0471-RSc0472).

### Two vertically inherited MCPs in the *Ralstonia* chemoreceptome contain a conserved motif for amino acid sensing

Recently, an amino-acid sensing motif of chemoreceptors was defined: YxxxxRxWY[x 13]Y[x 27-34]D (26). To search for the conserved amino acid binding motif in the *Ralstonia* MCPs, we aligned protein sequences of each MCP from GMI1000 and a known amino acid-sensing chemoreceptor: McfA (Pput_3489) from *P. putida* F1 (25, 27). We identified two chemoreceptor genes that contain the complete motif: RSc0606 and RSc3307 (Fig. 2A). RSc0606 was previously identified as an amino acid chemosensor (in *R. pseudosolanacearum* MAFF106611) and was named McpA (16). Hereafter, we call RSc0606 “McpA1” and RSc3307 “McpA2”. McpA1 and McpA2 share 54% amino acid identity with each other. Full-length *mcpA1* and *mcpA2* are present in all genomes except those of the genome-reduced phylotype IV strains that cause Sumatra Disease of Clove (SDC): R24 and NCPPB3219 (Fig. 1A and 2B). In the SDC lineage, the *mcpA1* and *mcpA2* genes are truncated by putative loss-of-function mutations.

**Figure 2.**
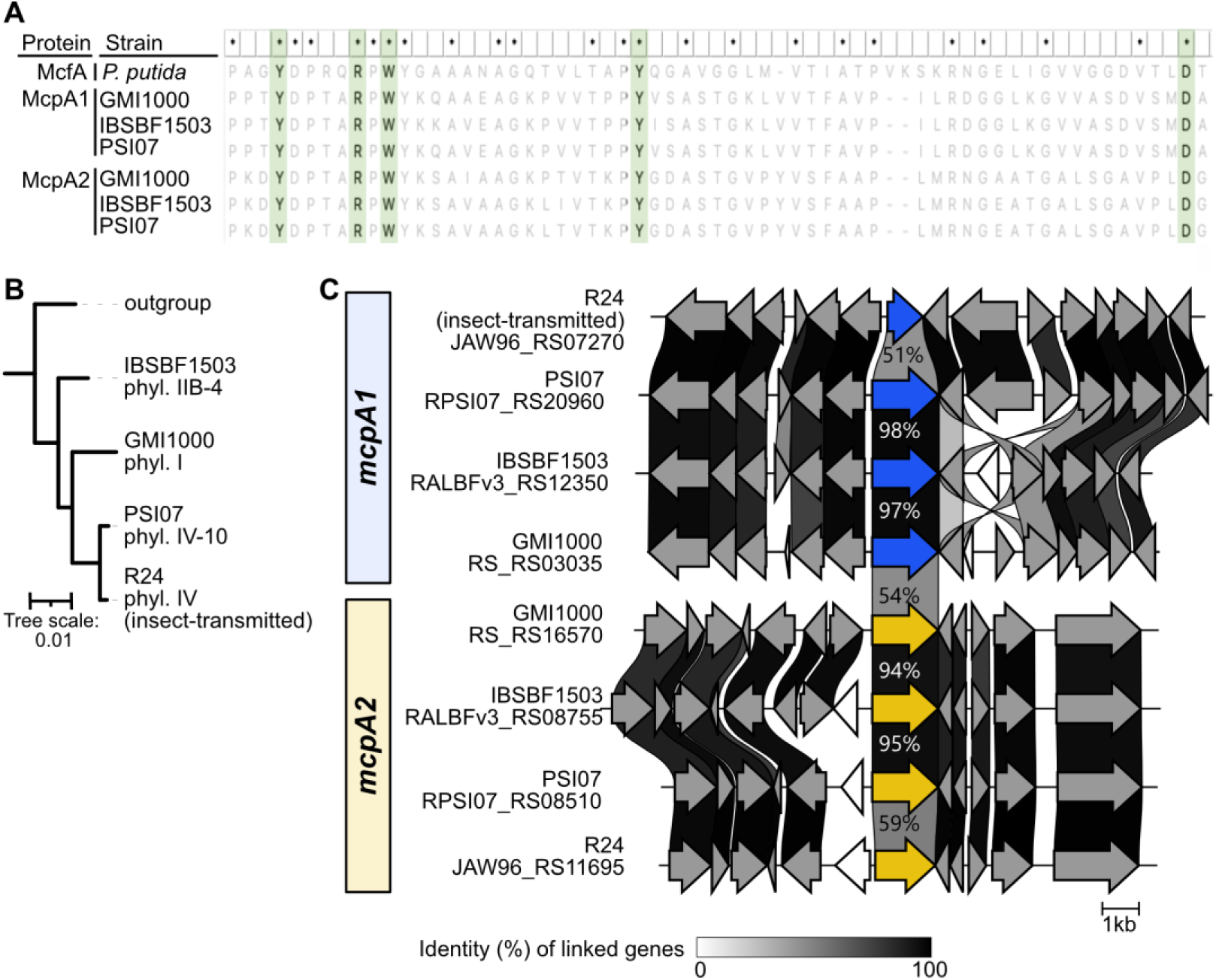
McpA1 and McpA2 contain an amino acid-binding motif, and the corresponding genes are vertically inherited within the *Ralstonia* wilt pathogens. (A) A multisequence alignment of the ligand binding domains of McpA1 and McpA2 shows the amino-acid recognizing motif: YxxxxRxWY[x13]Y[x27-34]D. McfA is included as a reference of a known MCP that binds amino acids. The MUSCLE alignment was created in MEGA v. 11. Green rectangles highlight the conserved motif residues. (B) A phylogenetic tree of the four genomes analyzed (GMI1000, IBSBF1503, PSI07, and R24) and an outgroup (*Ralstonia mannitolilytica*). The tree was built with the KBase SpeciesTree app. (C) Synteny of the gene neighborhoods for *mcpA1* (blue) and *mcpA2* (yellow) in the representatives genomes was visualized using Clinker (73). Homologous genes are connected by linkages that reflect the global amino acid identity of the pairings. The interactive HTML file from the Clinker analysis is available at doi.org/10.6084/m9.figshare.22337059.

### Chimeric HyChemosensors McpA1-NarQ and McpA2-NarQ are responsive to amino acids

To test whether the McpA1 and McpA2 chemoreceptors bind to amino acids as predicted by the bioinformatic model, we generated HyChemosensors (hybrid chemosensor-two component systems) (25) to probe the ligand-receptor interactions using a transcriptional reporter system in *E. coli*. To generate each HyChemosensor, we individually fused the N-terminal ligand binding regions of McpA1 and McpA2 to the HAMP domain and signaling domain of NarQ, the sensor kinase of a nitrate-responsive two-component system in *E. coli* that activates *narG* transcription (Fig. S6) (28–30). We constitutively expressed these MCP-NarQ hybrids from pHG165 in VJS5054 (30), an *E. coli* reporter strain that contains a genomic *narG* promoter::*lacZ* fusion (25). Previously, we reported that when individual attractant chemicals bind to an MCP-NarQ HyChemosensor, expression of the reporter protein decreases (25). This is due to the fact that a ligand-bound MCP naturally reduces the autophosphorylation activity of CheA. In contrast, when NarQ is bound to nitrate, it activates its own phosphorylation to trigger the activation of nitrate-responsive promoters. Therefore, a ligand-bound MCP-NarQ HyChemosensor reduces the autophosphorylation activity of the NarQ signaling domain. Thus, there is lower expression of the transcriptional reporter when a ligand is present.

First, we tested the McpA1-NarQ and McpA2-NarQ HyChemosensors against a complex mixture of amino acids, casamino acids (Fig. 3A). As a positive control for amino acid sensing, we used the *P. putida* McfA-NarQ HyChemosensor (25). All three HyChemosensors (McpA1-NarQ, McpA2-NarQ, and McfA-NarQ) exhibited reduced reporter expression when cultured in the presence of 1% w/v casamino acids. Although we have previously shown that HyChemosensors respond to individual ligands (25), this was the first time a HyChemosensor has been shown to detect a signal from a complex mixture of potential ligands.

**Figure 3.**
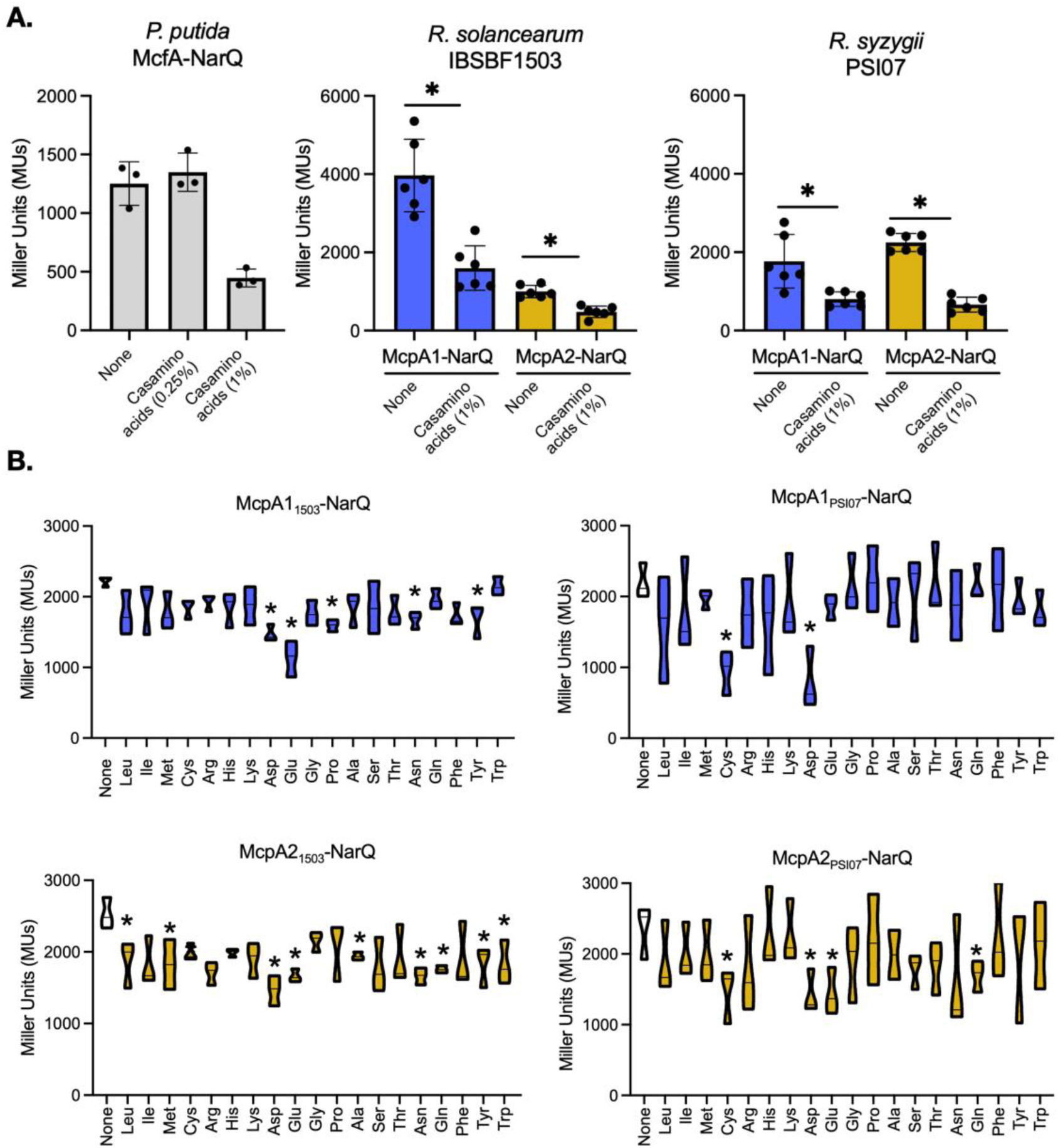
McpA1 and McpA2 HyChemosensors respond to amino acids. (A-B) As previously reported (25) and shown in Fig. S6, HyChemosensors have lower β-galactosidase reporter output when interacting with a ligand. Reporter strains were all in *E. coli* VJS5054 background and grown in MSB minimal medium with 80 mM glucose as a carbon source. Activity is reported in Miller Units (MUs). McpA1 and McpA2 orthologs were tested from two strain backgrounds: *R. solanacearum* IBSBF1503 (or “1503”) and *R. syzygii* PSI07. (A) Responses were measured for McfR-NarQ from *Pseudomonas putida* F1 (gray bars), McpA1-NarQ (blue bars), and McpA2-NarQ (yellow bars) with no attractant added (none), 1% w/v casamino acids, 0.25% w/v (McfR-NarQ only), or 5 mM Asn (McfR-NarQ only). Results show the averages from at least three independent experiments, and error bars represent standard deviation. (B) McpA1 and McpA2 have overlapping but distinct binding profiles for proteinogenic amino acids. All amino acids were added at 1 mM. Valine was excluded because it arrested growth of *E. coli* VJS5054 in MSB medium. Results are shown as box plots representing the median and range from at least three independent experiments. Asterisks indicate significance of amino acid treatment relative to the glucose only control by Student’s T-test, p<0.05.

Because the HyChemosensors can report a signal through the noise of a complex mixture, we attempted to eliminate specific amino acid ligands using amino acid subsets based on shared chemical characteristics (Fig. S7A). We screened the McpA1-NarQ and McpA2-NarQ using two orthologous periplasmic sensing domains from strains *R. solanacearum* IBSBF1503 and *R. syzygii* PSI07. From this point forward, we will refer to these HyChemosensors as McpA1_1503_-NarQ and McpA1_PSI07_-NarQ or McpA2_1503_-NarQ and McpA2_PSI07_-NarQ. The ligand binding regions of McpA1_1503_-NarQ and McpA1_PSI07_-NarQ have 95% amino acid identity and the ligand binding regions of McpA2_1503_-NarQ and McpA2_PSI07_-NarQ have 96% amino acid identity. Interestingly, despite the conservation of the binding regions, the McpA1 and the McpA2 from different strains displayed variable binding profiles across the amino acid pools (Fig. S7). McpA1_1503_-NarQ responded to four of the six pools containing 12 amino acids, while McpA1_PSI07_-NarQ responded to only three of the six pools. McpA2_1503_-NarQ responded to three of the six pools, while McpA2_PSI07_-NarQ responded to five of the six pools. To understand the specificity of McpA1 and McpA2, we then probed all four HyChemosensors against the individual proteinogenic amino acids (Fig. 3B). All four HyChemosensors had some level of binding to the acidic amino acids Asp and Glu. McpA1_1503_-NarQ also responded to Pro and Asn. McpA2_1503_-NarQ responded broadly to Ile, Met, Cys, Ala, Asn, Gln, Trp, and Tyr, whereas McpA2_PSI07_-NarQ recognized Cys and Gln.

### Mutants lacking *mcpA1, mcpA2*, or both have reduced responses to amino acids

We investigated the *in vivo* contributions of McpA1 and McpA2 to *Ralstonia* chemotaxis to amino acids. When creating chemoreceptor deletions in IBSBF1503 and PSI07 proved to be challenging, we constructed single deletions and a *mcpA1 mcpA2* double deletion in *R. pseudosolanacearum* GMI1000 (phylotype I). McpA1’s specificity for amino acids was previously demonstrated in *R. pseudosolanacearum* MAFF106611 (also phylotype I). McpA1 proteins from MAFF106611 and from GMI1000 have 100% amino acid identity. We continued with GMI1000 as a model phylotype I strain to allow us to directly compare to the previously reported characteristics of McpA1. To probe the functional contributions of McpA1 and McpA2 in *Ralstonia*’s response to amino acids, we used quantitative swim plate assays.

Quantitative swim plate assays require strains to catabolize the target attractant (31). As the attractant is catabolized, a chemical gradient is produced that guides motile cells to swim outward from the point of inoculation. As expected, the Δ*mcpA1* and Δ*mcpA2* single mutants and the Δ*mcpA1*Δ*mcpA2* double mutant were significantly less attracted to casamino acids (Fig. S8). The responses of the single mutants and the double mutant were not statistically distinct from one another (Student’s T-test: Δ*mcpA1*Δ*mcpA2* versus *ΔmcpA1* p=0.20; versus Δ*mcpA2* p=0.19).

Next, we probed the response of *Ralstonia* wilt pathogens to individual amino acids. To account for the natural diversity of these pathogens, we tested chemoattraction across three model strains: IBSBF1503, PSI07, and GMI1000. Because soft agar assays require catabolism of the target attractant, we tested the ability of the *Ralstonia* model strains to use each of the proteinogenic amino acids as sole sources of carbon and energy. All three strains had similar carbon utilization patterns and were capable of using 11 of the 20 proteinogenic amino acids as sole sources of carbon: Ala, Asn, Asp, Glu, Gln, His, Pro, Ser, Thr, Trp, and Tyr (Fig. S9). Of the eleven carbon sources, all three strains displayed a chemotactic response in swim plates to only five: Asp, Asn, Glu, Gln, and Ala (Fig. S9). Soft agar plates containing 1 mM His, Ser, Thr, and Trp showed clear growth at the inoculation site, but no characteristic halo formation after 4 days of incubation. Only PSI07 showed attraction to Tyr in soft agar plates while GMI1000 and IBSBF1503 did not. In addition, only GMI1000 responded to proline.

To quantitatively study chemoreception and response, model bacterial strains are often serial passaged to enrich for highly motile cell populations (31). Enriched strains out-perform their parental environmental isolates in swimming motility assays (32, 33). Because enrichment results in a stronger chemotactic response, swim plate experiments can provide clear results because assays can yield clear results before evaporation has increased the rigidity of the agar. To differentiate the subtle differences in chemotactic responses of *Ralstonia mcp* mutants, we enriched the three strains for the swimming phenotype by serial passaging. For each strain, the initially isogenic population was grown on soft agar CPG plates and the population at the fringes of the swarmed halo was serially passaged on fresh soft agar plates. After three passages, we isolated single colonies that we designated SWM1000, SWM1503, or SWM07 based on the strain background (Fig. S10A). Wildtype *Ralstonia* have a slow chemotactic response in the rich CPG medium, taking up to 48 hours to generate a measurable halo. In contrast, SWM1000 generated halos roughly 4 times as large as those generated by the parental GMI1000 wild type after 16-hr incubations at 28 °C (Fig. S10B). The increased halo size of SWM07 and SWM1503 were less pronounced than SWM1000. To ensure that the rapid development of the chemotactic halo was not due to an increased rate of growth, we compared the growth of SWM1000 and GMI1000 in liquid CPG and in minimal medium with selected amino acids as carbon sources. We observed no difference in growth phenotype between GMI1000 and SWM1000 (Fig. S11). Growth was also compared for SWM1503 and SWM07 to their respective wildtype parents, IBSBF1503 and PSI07, and similarly, we found no growth phenotype between the swarm strains and the parental wild types (data not shown).

Taking advantage of the improved response time of the enriched isolate, SWM1000, we quantified this strains’ response to individual amino acids in minimal medium (Fig. 4). SWM1000 Δ*mcpA1,* SWM1000 Δ*mcpA2,* and the SWM1000 Δ*mcpA1* Δ*mcpA2* double mutant were significantly less attracted to Asp, Glu, and Pro. The loss of McpA2 subtly reduced the SWM1000 response to Asn, while the loss of both receptors did not significantly reduce SWM1000 attraction to Gln and Ala (Fig. 4). Further, the loss of *mcpA1* resulted in a reduced ability to swim towards Gln that was not observed in the strains lacking *mcpA2* (Fig. 4).

**Figure 4.**
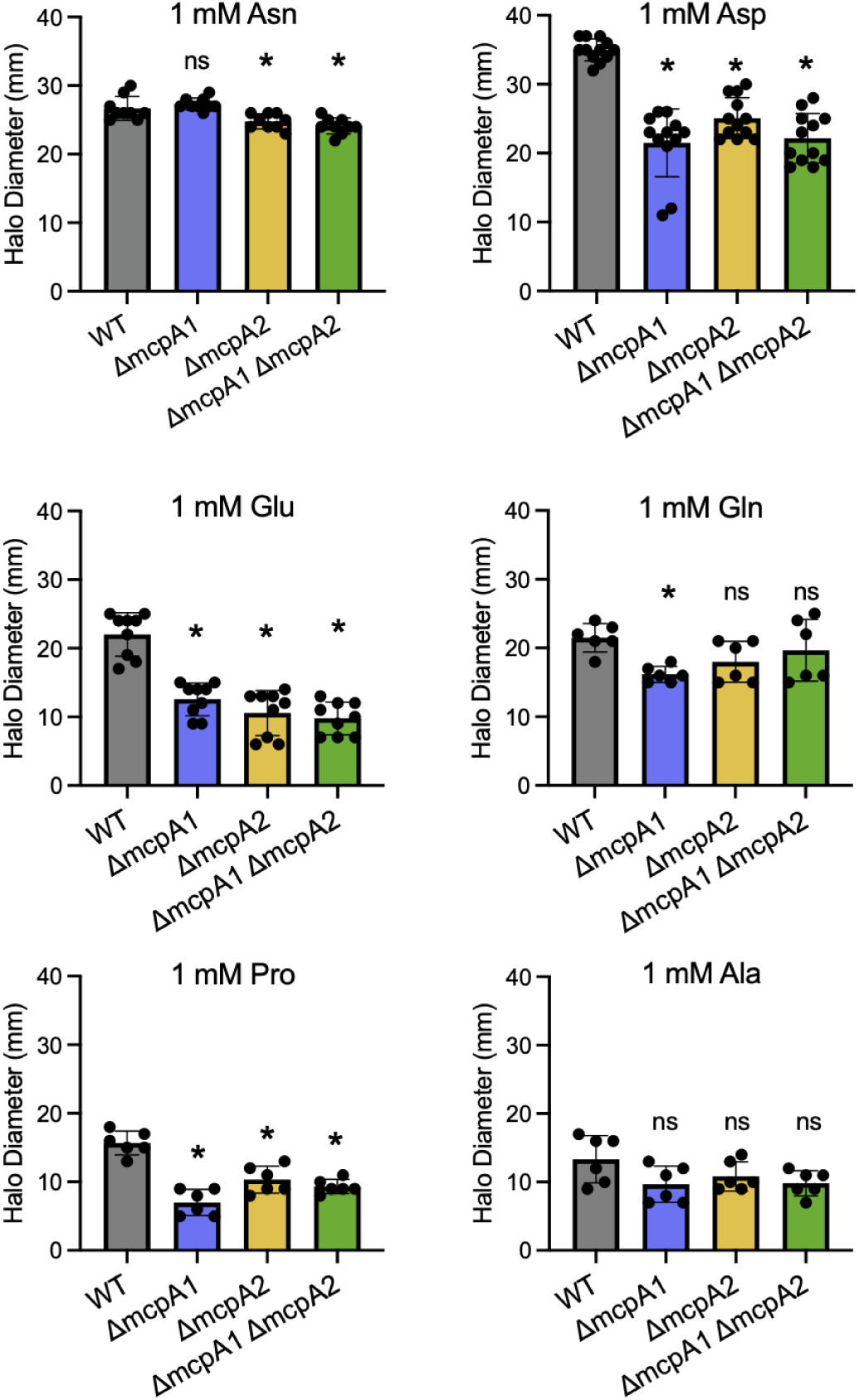
McpA1 and McpA2 contribute to *Ralstonia* chemotaxis for individual proteinogenic amino acids. All strains were constructed in the SWM1000 background and include wildtype (WT), Δ*mcpA1*, Δ*mcpA2*, or Δ*mcpA1* Δ*mcpA2* genotypes. Strains were inoculated into minimal MSB medium containing 0.3% w/v agar and 1 mM of the indicated amino acid. The diameter of each halo was measured after incubation at 28°C after 50-60 hr. Data points represent at least two trials each with three biological replicates per trial. Asterisks indicate significance relative to the wild type using a Student’s T-test control (* = p < 0.05; ns = not significant).

### McpA2 contributes to *R. pseudosolanacearum*’s ability to infect tomato plants after soil inoculation

To understand the biological role of McpA1 and McpA2 in *Ralstonia* infections, we challenged our single and double mutants to infect tomato via soil soak inoculation. Because the previous *mcpA1* study used phylotype I *R. pseudosolanaceaum* strains (MAFF106611 and Ps29), we used phylotype I strain GMI1000. In agreement with Hida *et al.* 2015, we observed that infection with GMI1000 Δ*mcpA1* was not significantly different from that of wild type GMI1000. In contrast, both GMI1000 Δ*mcpA2* and GMI1000 Δ*mcpA1* Δ*mcpA2* showed a trend of reduced virulence after soil drench inoculation (Fig. 5A). To robustly analyze virulence defects, we also tested virulence in the SWM1000 background by challenging SWM1000 Δ*mcpA1* and SWM1000 Δ*mcpA2,* and double mutant SWM1000 Δ*mcpA1* Δ*mcpA2* to infect tomato using naturalistic soil soak inoculation. In this strain background, McpA2 was clearly required for full virulence while McpA1 remained dispensable for virulence (Fig. 5B). Notably, when we inoculated strains directly into the stem of tomato using a cut petiole inoculation method, neither the loss of McpA1 or McpA2 affected the rate of disease progression (Fig. S12). Thus, McpA2 contributes to invasion but not replication in the plant.

**Figure 5.**
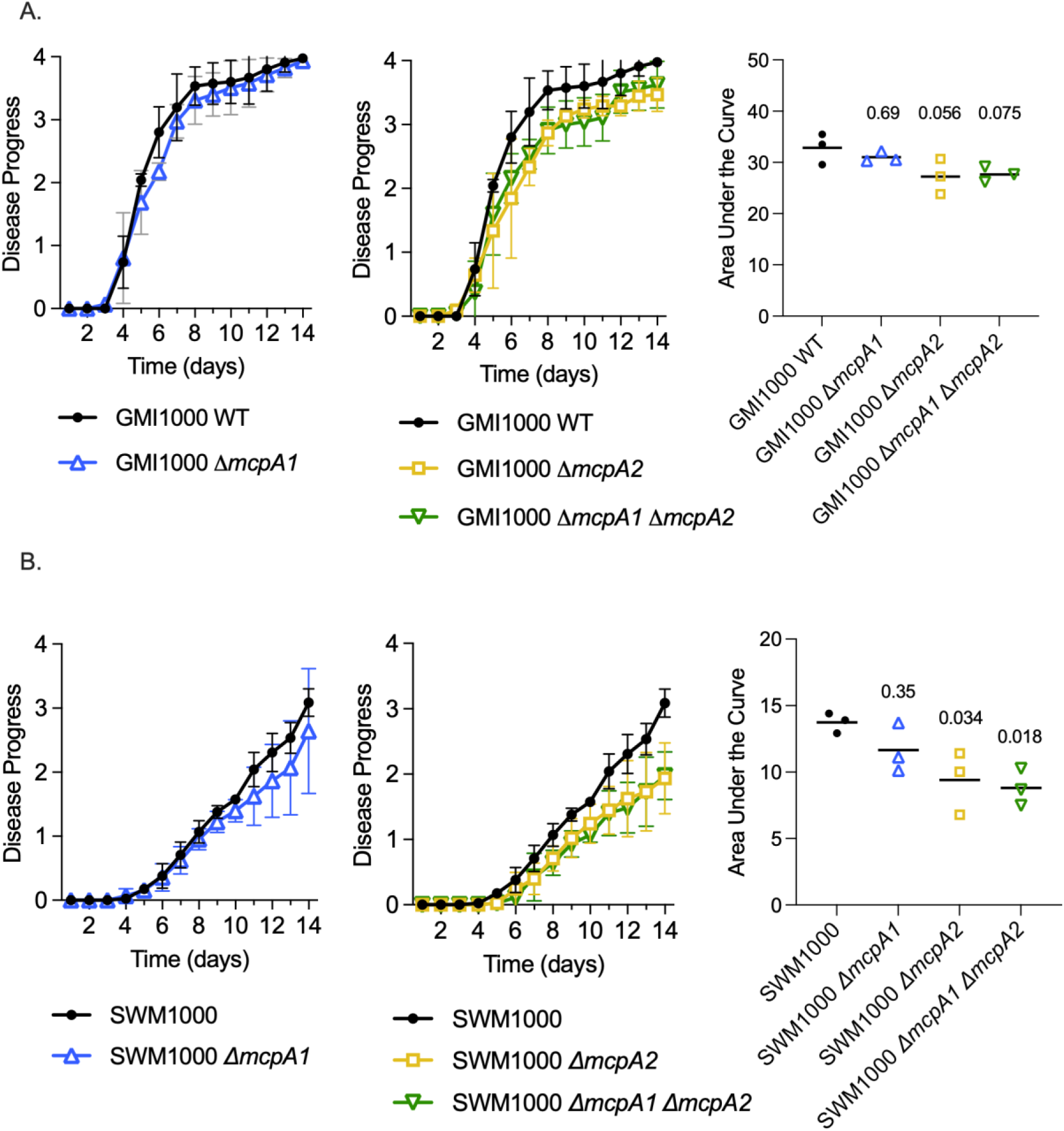
*mcpA2* contributes to *Ralstonia* virulence after soil drench inoculation. Tomato cv. moneymaker plants were inoculated by pouring wildtype (WT), Δ*mcpA1*, Δ*mcpA2*, or Δ*mcpA1* Δ*mcpA2* strains in a GMI1000 background (A) or a SWM1000 background (B) onto the soil. Plants were rated daily on a disease index scale from 0 to 4 for 14 days. Each data point is the average disease index for that day across three independent trials (n=15 plants per trial). The same trial data is also represented for both (A) GMI1000 background and (B) SWM1000 background as Area Under the Curve (AUC). P-values reflect results of a Ordinary One-Way ANOVA in comparison to the WT control (either SWM1000 or GMI1000).

### SWM1000 is less virulent than GMI1000 in naturalistic soil soak assays

Although SWM1000 was capable of causing wilt disease on tomato, symptom onset was delayed compared to GMI1000 (Fig. 5). Because motility and biofilm production are co-regulated in many bacteria (34, 35), we quantified the biofilm production of SWM1000 and GMI1000 using a PVC biofilm assay. However, the two strains generated equivalent biofilms in the *in vitro* assay (Fig. S13). A previously studied hyper-motile mutant of *Ralstonia* strain K60 (Δ*motN*) out-performed wildtype isolates of *Ralstonia* in swimming motility assays in a metabolism-independent manner, but also exhibited reduced virulence in soil soak assays (36). Unlike the *motN* mutant, we observed SWM1000 to have pleiotropic phenotypes. For example, the SWM1000 displayed increased production of diffusible melanin pigments in addition to the selected motility trait. Although increased melanin production is a phenotype of *phcA* quorum sensing mutants (37), the SWM1000 mutant produces mucoid colonies, indicating its quorum sensing system is intact. Because SWM1000 was isolated following serial inoculations and had genetically stable phenotypes, we sequenced the genome of SWM1000 to identify underlying mutations. We compared the sequences of SWM1000 and GMI1000 using BreSeq, a computational pipeline for identifying sequence variation from short read sequences (38). Mutations in *phcA, motN*, or other chemotaxis or flagellar genes were not observed. The pipeline identified mutations in several IS elements (Table S5), which may perturb gene expression in the strain. Due to the repetitive nature of IS elements in genomes, it is not clear which, if any, of these are causative of the SWM1000 phenotype(s).

## Discussion

It is not unusual for MCPs in metabolically versatile organisms to overlap in their chemical binding profiles (23, 24). Here, we identify that in addition to McpA1 (previously “McpA” (16)), *Ralstonia* have a second chemoreceptor, McpA2, that redundantly detects amino acids. HyChemosensor reporters and behavioral assays showed each ortholog of McpA1 and McpA2 responds to acidic amino acids, Asp and Glu, as well as a handful of other amino acids. Notably, the McpA2_1503_-NarQ and McpA2_PSI07_-NarQ HyChemosensors responded to a wider range of amino acid ligands than McpA1 HyChemosensors, indicating functional differences between the McpA2 orthologs.

When challenged to infect a host by naturalistic soil soak inoculations, McpA1 and McpA2 did not contribute equally to *Ralstonia* physiology. Our results corroborate the prior genetic studies showing that loss of McpA1 is not sufficient to influence *Ralstonia’*s host detection (16). If MpcA1 is contributing to amino acid sensing during host detection, the activity is either redundant with other MCPs or masked by an energy taxis receptor. Due to the functional overlap HyChemosensor assays and the additive contributions to chemotaxis to casamino acids in swim plate assays, we expected that McpA2 would be functionally redundant with McpA1. However, we found the loss of McpA2 has a dominant influence on host detection. The loss of McpA2 resulted in a significant defect in infection by naturalistic soil soak, even when the McpA1 protein was present. These results indicate it is unlikely that these two MCPs are simply redundant, but rather McpA2 plays a physiologically distinct role in locating and colonizing the host from the soil. Importantly, virulence of Δ*mcpA2* and Δ*mcpA1* Δ*mcpA2* mutants were not distinct from that of wild type when these mutants were inoculated by cut petiole inoculations, showing that the ability to detect amino acids is critical when the bacterium locates the host from the soil, not for virulence within the host.

### Amino acids commonly serve as signals for host-bacterial interactions

The amino acid receptor McpA2 joins the malate receptor McpM as being necessary for *Ralstonia*’s full virulence following soil soak (16). These receptors presumably allow *Ralstonia* to locate entry points in plant roots. Roots exude up to 40% of the plant’s fixed carbon as a milieu of organic molecules (39). For example, tomato root exudates contain many simple metabolites: over 52 unique sugars, 14 sugar alcohols, 25 amino acids, 5 fatty acids, 39 organic acids, and 5 simple phenolics in addition to over 100 complex metabolites, including 9 hydroxycinnamate conjugates, 33 acylsugars, and 5 alkaloids (40, 41). Root exudation can be affected by biotic and abiotic stresses (42–46); notably in abiotically-stressed plants, amino acid levels in exudates are decreased (45). Moreover, as *Arabidopsis* plants age, sugar exudation decreases while amino acid or phenolic compound exudation increases (47). This variation in root exudate chemistry likely fine-tunes the attractiveness of roots to soil bacteria.

The ubiquity of amino acids within plant exudates might make them broad signals that alert phytobacteria regarding the presence of a potential host. Like *Ralstonia*’s McpA1 and McpA2, phytopathogenic *P. syringae* virulence requires chemotactic responses to Asp and Glu (48). Loss of the L-Asp/D-Asp/L-Glu receptor PscA reduced the virulence of *P. syringae* on tomato. Notably, exogenous L-Asp reduced *P. syringae* virulence, indicating that endogenous Asp gradients contribute this pathogen’s ability to invade hosts (48). *Pectobacterium atrosepticum,* a soil borne pathogen of potato, has three overlapping amino acid receptors, PctA, PctB, and PctC, that allow it to detect 19 of the 20 proteinogenic amino acids (22). *Dickeya dadantii* also shows a strong chemotactic response to Asp, Glu, and other amino acids, but specific receptors have not yet been characterized (49). Non-pathogen phytobacteria including *Sinorhizobium meliloti* (McpU (50))*, Bradyrhizobium japonicum* (51), and *Azotobacter chroococcum* (52) are also capable of sensing these amino acid signals and responding to plant root exudates.

The versatility of amino acids as a host signal extends beyond bacterial phytopathogens: amino acid sensing-MCPs also critical in host-microbe interactions for several animal pathogens. *P. aeruginosa, V. cholera, H. pylori*, *S. enterica*, and *E. coli* require chemotaxis for full virulence (Reviewed in (53)). Animal pathogens tend to have fewer MCPs encoded in their genomes than environmental bacteria; however, a notable number of chemoreceptors within these organisms detect amino acids. *P. aeruginosa* has 3 amino acid receptors: PctA (Trp/Met/Ile), PctB (Arg/Gln), and PctC (GABA) (54). Wildtype *P. aeruginosa* migrates to wounded epithelial cells using chemotaxis: notably the loss of all three of these amino acid receptors was required to generate a mutant incapable of detecting the wounded host cells (24). Pathogenicity of *V. cholera* was found to be significantly reduced when McpX (Ser/Arg/Asp) was removed (55). *E. coli* has four MCPs (Tsr, Tar, Trg, and Tap), and two of those MCPs (Tsr and Tar) detect amino acid ligands (56). Interestingly, amino-acid-sensing Tsr and Tar are generally absent within uropathogenic *E. coli,* while enteric and commensal *E. coli* tend to have all four receptors (57), showing that the life style of the chemotactic organism influences the retention of *mcp* genes. All of these examples, show that amino acid detection plays a role in host sensing across plant and animal hosts.

### Loss of MCPs in an insect-vectored *Ralstonia* lineage

Despite the overall conservation of MCPs in the *Ralstonia* pangenome, there is a notable exception. The otherwise conserved McpA1 and McpA2 are truncated as pseudogenes in genomes of phylotype IV strains that cause Sumatra Disease of Clove (SDC): R24 and NCPPB3219. The SDC pathogens are unusual members of the *Ralstonia* wilt pathogens. Instead of having a soil-borne life cycle, SDC pathogens are vectored by xylem-feeding insects (58). Consistent with the lifestyle switch, genomes of SDC strains have signs of genome reduction (58). The loss of functional forms of the McpA1 and McpA2 receptors supports the model that chemotaxis towards amino acids contributes to *Ralstonia*’s ability to locate and invade roots.

Because the SDC lineage has this unique ecology and loss of functional McpA1 or McpA2, it begs the question of whether we can identify additional target MCPs from these genomes as critical for soil vectored transmission of *Ralstonia.* In addition to pseudogenized *mcpA1* and *mcpA2*, the SDC lineage lacks the *aer1* energy taxis chemoreceptor. This energy taxis gene was also truncated in a soil-borne lineage of phylotype IV containing strains T95, T51, SL2064, and KACC10722. The SDC strains have maintained the remainder of the core MCPs (Fig. 2). The other core *Ralstonia* receptors are well-conserved across the phylotypes, indicating that they are important enough to *Ralstonia* physiology to persist, or at least not costly enough to be maintained in the genome. Each is potentially involved in the ability to sense additional root exudates. To date, *Ralstonia* are known to respond to diverse root exudate chemicals from sugars and organic acids, to phytohormones like salicylic acid (4, 59). Importantly, other characterized MCPs sense root exudate compounds (e.g., citrate, malate), but with the exception of *mcpM,* the loss of the MCP does not cause a virulence defect (16–18, 60).

### The chemoreceptor repertoire is more consistent than repertoires of secreted effectors

*Ralstonia* pathogens have high genome plasticity, enabling their adaptability to adapt in the face of dynamic ecological stresses. Here, we investigated the pangenome plasticity of the previously identified large repertoire of *Ralstonia* chemotaxis receptors (16). Orthologous MCPs were highly conserved in the *Ralstonia* pangenome, suggesting that there is not strong selection to diversify chemotaxis specificity by gain and loss of MCPs. This is in contrast to the dynamic gain-and-loss patterns of plant-manipulating type 3 secreted effectors and microbiota-fighting type 6 secreted toxins (61, 62). The MCPs were evenly distributed across the chromosome and megaplasmid, unlike the observed megaplasmid-enrichment of type 3 effectors and type 6 toxins (61, 62). Despite dynamic gene flow on the megaplasmid (63), ten MCPs and the majority of the chemotaxis and flagellar machinery were well-conserved on the megaplasmid. Future studies should investigate the phenotypic plasticity conferred by sequence variation in the MCPs and the presence of the rare accessory MCPs.

### Enhanced motility does not accelerate bacterial wilt disease

Chemotaxis is a well-documented virulence factor for many phytopathogens (2, 3, 53), but *Ralstonia* strains with increased motility do not show increased virulence (36). SWM1000-background strains responded more rapidly to chemical gradients in a growth-independent manner but had a delayed ability to infect the host from the soil.

It is interesting that strains with increased motility or chemotaxis phenotypes are worse at locating a host from the soil. Why is this the case? One hypothesis is that increased motility interferes with the cell’s ability to navigate its environment (64, 65). Increased responses to steep chemical gradients may come at a trade-off to detecting subtle chemical gradients (65). In this scenario, increased motility and chemotaxis could lead to the cell swimming past the root. Another hypermotile *Ralstonia* mutant, with a defect in *motN,* shares a similar virulence phenotype to SWM1000 (36). MotN is involved in regulating the flagellar-mediated motility phenotype in *Ralstonia*. The previously characterized *motN* mutant is hyperflagellated and has increased motility, altered biofilm formation, and reduced virulence on tomato via naturalistic soil soak. Like the *motN* mutant, SWM1000 has reduced virulence only when infecting from the soil, but SWM1000 does not have any mutations within or around the *motN* gene nor any changes in its biofilm formation phenotype. The mutations that cause increased motility in SWM1000 are unclear, but BreSeq analysis revealed several point mutations in IS element transposases relative to the wildtype genome. Alteration of one or more of IS elements could result in a regulatory shift in bacteria (66). Additional investigation of enriched isolates like SWM1000 may reveal more information about how motile behavior is regulated in bacterial plant pathogens with complex life cycles like *Ralstonia*.

## Materials and Methods

### Bacterial Growth Conditions

Bacterial strains and plasmids that are used in this study are included in Table S2 and S3. *E. coli* strains were cultured in Luria-Bertani (LB) medium (10 g/L tryptone, 5 g/L yeast extract, and 5 g/L NaCl) incubated at 37°C. *Ralstonia* strains were grown in Casamino Acids-Peptone-Glucose (CPG) broth media (1 g/L casamino acids, 10 g/L bacto-peptone, 5 g/L glucose and 1 g/L yeast extract) at 28°C. To monitor that *Ralstonia* maintains virulence, solid CPG media contains 0.5% (w/v) 2,3,5-Triphenyltetrazolium chloride (TZC) unless otherwise noted. TZC is a colorimetric media additive that monitors redox activity through the accumulation of a red pigment. Altered colony morphology or increased accumulation of red pigmentation allows for the exclusion potential of spontaneous avirulent mutants from further cultivation and experimentation. When defined minimal medium was required, *E. coli* and *Ralstonia* strains were cultivated in modified Hutner’s Mineral Base (MSB (67)) medium (25 mM potassium phosphate buffer pH =7.3, 755 mM (NH_4_)_2_SO_4_, and 10.5 mM nitrilotriacetic acid (NTA-free acid), 25 mM KOH, 23 mM MgSO_4_, 4.5 mM CaCl_2_.2H_2_O, 1.4 μM (NH_4_)_6_Mo_7_O_24_.4H_2_O, 8.5 μM EDTA, 38 μM ZnSO_4_•7H_2_O, 87 μM FeSO_4_•7H_2_O, 9 μM MnSO_4_•H_2_O, 1.5 μM CuSO_4_•5H_2_O, 800 nM Co(NO_3_)_2_•6H_2_O, and 460 nM Na_2_ B_4_O_7_•10H_2_O). Amino acids were added to MSB liquid medium at a final concentration of 1 mM or 5 mM. Antibiotics were added to the medium when required for selection, screening, or plasmid maintenance: kanamycin at 25 mg/L for *Ralstonia* and 50 mg/L for *E. coli* and ampicillin at 100 mg/L.

### Identifying MCPs in the Ralstonia pangenome

We first used the JGI Integrated Microbial Genomics (IMG) platform (68) to identify all MCPs in the genomes of species reference strains (GMI1000, IBSBF1503, and PSI07). The motif for the MCP cytoplasmic signaling domain (pfam000015) was queried against the genomes using the Function Search tool. We characterized the domain architecture of each MCP family using the InterProScan Tool from EMBL (69).

We analyzed MCP gene presence/absence in a set of high-quality, publicly available *Ralstonia* genomes previously defined in (62). The initial goal was to analyze complete genomes, but this would have excluded almost all phylotype II, III, and IV genomes. Thus, we selected genomes assembled into fewer than 28 contigs with CheckM completeness > 99.82% and CheckM contamination < 0.96% (70).

In KBase (71), we used the Build SpeciesTree app to generate a phylogenetic tree based on 49 conserved genes. The .newick tree file was uploaded into iToL for visualization (72). We queried each GMI1000 MCP sequence against the genomes using the BLAStP app in KBase with identity > 50%, coverage > 80% thresholds. The percent identity of the BLAST hits for each MCP query revealed clear thresholds that allowed us to manually define orthologs. The presence/absence of each MCP gene family was visualized on iToL using the iToL annotation editor.

To infer whether MCP genes are vertically inherited, we used Clinker (73) to perform a synteny analysis. From the NCBI Graphics View, we downloaded GenBank Flat files of the genetic neighborhood surrounding each *mcp* gene in GMI1000, IBSBF1503, and PSI07. Then, we converted the files to .gbk format and analyzed and visualized the synteny with Clinker. Analysis of these genetic neighborhoods provided an independent validation that we had correctly defined orthologous MCPs based on BLAST identity.

### Strain Construction and Mutagenesis

Standard serial passage was used to isolate hypermotile “swarm” strains in the background of *R. pseudosolanacearum* GMI1000, *R. syzygii* PSI07, and *R. solanacearum* IBSBF1503: SWM1000, SWM07, and SWM1503, respectively. Each wildtype strain was inoculated onto CPG agar without TZC and grown at 28°C for 2 days. A single colony was inoculated into the center of a soft agar plate containing 50% v/v CPG and 0.25% w/v agar. After 2 days of incubation at 28°C, the inoculated bacteria had produced a typical chemotaxis halo (31). The exterior border of the halo was scraped with a sterile stick and inoculated into a fresh soft agar plate. This serial passage enrichment was repeated for a total of 3 serial passages. To isolate SWM1000, SWM07, and SWM1503, the halo from the fourth soft agar passage was streaked to CPG and re-streaked to CPG containing 1% TZC. Single colonies of the swarm mutant and the ancestral wild type were inoculated into CPG containing 0.25% agar to qualitatively confirm that swarm strains had stronger chemotactic responses in soft agar. GMI1000 and SWM1000 were sequenced using Illumina Short Read amplification and assembled in KBase, and annotated by Prokka (74). Breseq (38) was used to identify variations in the short read data between GMI1000 and SWM1000.

<u>Construction of deletion mutations:</u> All primer sequences used for the generation of deletion mutations are listed in Table S4. The Δ*mcpA1* and Δ*mcpA2* markerless deletions were created through a selection with kanamycin and *sacB*-dependent counter-selection with pKD46-derivative plasmids. Specifically, we used Gibson assembly to create the knockout plasmids, pJKA002 and pJKA005. For pJKA002: The primers JKA_22 and JKA_23 were used to amplify the 1159 bp upstream of *mcpA1.* The primers JKA_24 and JKA_25 were used to amplify the 760 bp downstream of *mcpA1*. These primers were designed to generate amplicons containing 5’ overlap regions with each other and the pKD46 plasmid at restriction sites for HindIII and BamHI using the NEBuilder software. The two amplicons and linearized vector were assembled using NEB HiFi Assembly Master Mix and transformed into *E. coli* DH5a using LB with kanamycin as the selective medium. The resulting construct was confirmed using restriction digest confirmation and Nanopore sequencing. The same protocol was used for pJKA005, except the primers JKA_18 and JKA_19 were used to amplify the 1105 bp upstream of *mcpA2* and JKA_20 and JKA_21 were used to amplify the 1115 bp downstream of *mcpA2*.

Electroporation was used to transform GMI1000 and SWM1000 with the plasmids pJKA002 and pJKA005. Selection on CPG with kanamycin was used to select for integration of the plasmid into the *Ralstonia* genome. We then subjected transformants to a growth selection on 5% w/v sucrose and screened for kanamycin sensitive and sucrose resistant colonies. pJKA005 was also used to transform JKA008 (SMW1000 Δ*mcpA1*) and JKA004 (GMI1000 Δ*mcpA1*) by electroporation to generate the double mutation strains: JKA010 (SWM1000 Δ*mcpA1* Δ*mcpA2*) and JKA007 (GMI1000 Δ*mcpA1* Δ*mcpA2*). All strains were screened by PCR amplification to confirm deletion of the gene and for the possibility of a merodiploid construct using primers JKA_18 and JKA_25 from *mcpA1* deletions and primers JKA_18 and JKA_21 for *mcpA2* deletions.

### Quantitative Chemotaxis Assays in Soft Agar

Quantitative swim plate assays were adapted for *R. pseudosolanacearum* strains (31, 76). Strains were grown overnight (∼16 hr) in CPG and sub-cultured (1:50) into MSB containing 0.4% w/v glucose. Cultures were incubated at 28°C until mid-log cultures with OD_600_ of 0.3-0.4 were achieved. Bacteria were harvested by centrifugation at 5000 rcf and resuspended in MSB to an OD_600_ of 0.4 ± 0.02. Each well of an untreated 8-well rectangular plate (Nunc™ 267062) was filled with exactly 8 mL of MSB containing 0.25% w/v agar and either no ligand, 1% w/v casamino acids, or 1 mM of the test ligand. Three microliters of resuspended culture were inoculated into the center of each well. After incubation at 28°C for 16-to-20 hr, the diameter of the halo was measured to the nearest millimeter. Averages are the combination of at least three biological replicates for each condition. Responses to 1% w/v casamino acids and 50% v/v CPG were used as positive controls.

### Bio-reporter HyChemosensor assays

#### Construction of HyChemosensor strains

Bioreporter plasmids were generated by the DOE Joint Genome Institute. Briefly, DNA fragments encoding the ligand binding regions from the N-terminus through the established fusion site within the HAMP domain of McpA1 and McpA2 (RSc3307) from *R. syzygii* PSI07 and *R. solanacearum* IBSBF1503 were synthesized (25). The synthesized region was identified by MAFFT protein sequence alignments with NarQ and PcaY. pVJS3354, which contains the full length *narQ* gene, was linearized by NdeI digestion and Golden Gate assembly was used to fuse the synthesized ligand binding regions to the HAMP and signal transduction region of NarQ, generating pBSR21 (pVJS3354::mcpA_1503_-narQ), pBSR1 (pVJS3354::mcpA_PSI07_-narQ), pBSR16 (pVJS3354::RSc3307_1503_-narQ), and pBSR8 (pVJS3354::RSc3307_PSI07_-narQ). pBSR21, pBSR1, pBSR16, and pBSR8 were used to chemically transform *E. coli* VJS5054 (30) and transformants were selected on LB containing ampicillin.

#### Reporter assays

Reporter assays tracking β-galactosidase activity were performed as previously described (25). Briefly, because a low O_2_ concentration is required for *E. coli* to produce NarL, a critical component of the reporter system, we exploited microaerophilic growth conditions for the reporter assays. VJS5054 containing the HyChemosensor plasmids was inoculated into 5 mL LB medium and incubated with shaking at 37 °C for 16 hr. Strains were then sub-cultured 1:100 into 13 mL screw-cap tubes containing MSB medium with 80 mM glucose and 100 μg/mL ampicillin. To establish a microaerophilic environment, tubes were completely filled with medium to eliminate all of the headspace. Cultures were grown to mid-exponential phase (OD_600_ = 0.3-0.4), standing in a 37°C water bath incubator. Bacteria were harvested by centrifugation and resuspended in Z-buffer (0.1 M sodium phosphate pH 7.0, 10 mM KCl, 1 mM MgSO_4_, and 50 mM 1,4-dithiothreitol) to an OD_600_ of 0.4. For each assay, 1 mL of culture was solubilized in chloroform (0.1% v/v) and sodium dodecyl sulfate (17 mM). Activity was measured at room temperature by the addition of 0.2 mL of ortho-nitrophenyl-ꞵ-galactoside (4 mg/mL). Reactions were stopped by the addition of 0.5 mL of 1 M Na_2_CO_3_. As previously described, reporter output was expressed using arbitrary units, Miller Units (77). Each culture was assayed in triplicate and reported values are averages of 3 separate biological replicates.

#### Biofilm Assays

Biofilm assays were performed as previously described (3). *Ralstonia* strains were inoculated into CPG medium and grown for 16 hr at 28 °C. Cells were harvested by centrifugation and resuspended in fresh CPG medium. The bacterial suspension was adjusted to an OD of 0.1 or approximately 10^8^ CFU/mL. Polyvinyl chloride (PVC) 96-well plates were used as the biofilm formation surface. Each well was filled with 95 μl of fresh CPG medium and inoculated with 5 μl of the adjusted bacterial suspension. The plates were sealed with Breathe-Easy™ membranes (Sigma-Aldrich) to maintain sterility while allowing gas exchange. Plates were grown statically for 24 hr at 28°C. Following incubation, the wells were stained with 25 μl of 1% w/v crystal violet solution for 25 min at room temperature. Excess stain was removed by pipette and carefully washed twice with 200 μl of sterilized water.

To quantify biofilm formation, the adhered crystal violet was dissolved in 200 μl of 95% ethanol. The resulting solution was transferred to a new PVC plate, and absorbance was measured at 590 nm using a spectrophotometer. Uninoculated wells containing only CPG was run through the same method, and the resulting average of absorbance at 590 nm was subtracted as background. Higher absorbance values indicate greater biofilm formation.

#### Disease Progress Assays

Tomato cv. Moneymaker seeds were sown in Sunshine Mix #1 and grown in growth chambers maintained at 28°C with a 12-h light cycle. After 14 days, seedlings were transplanted to individual pots.

At 21-days-old, plants were inoculated. To prepare the inoculum, strains were grown on CPG agar containing TZC and incubated at 28°C for 48 hr. Overnight cultures were grown in CPG broth overnight at 28°C with shaking. For soil soak inoculations, tomato cv. Moneymaker plants were inoculated by slowly pouring 50 mL of bacterial suspension (OD_600nm_ of 0.1) around the plant stem, corresponding to 5 x 10^7^ to 1 x 10^8^ CFU/g soil. Plants were observed and rated daily for 14 days following inoculation. For cut petiole inoculations, the oldest true leaf was identified and cut horizontally using a sharp razor blade to expose a ∼2 mm petiole wound, and a 2 μL droplet of bacterial suspension containing ∼1000 total CFU was reverse pipetted directly to the wound. Plants were observed and rated daily for 14 days following inoculation. Disease index (DI) was measured using a standard disease index scale from 0 to 4, where 0 = 0% of leaves wilted, 1 = 0.1 to 25% of leaves wilted, 2 = 25.1 to 50% of leaves wilted, 3 = 50.1 to 75% of leaves wilted, and 4 = 75.1 to 100% of leaves wilted.

## Data Availability

Synteny analyses and input files are available on FigShare at doi.org/10.6084/m9.figshare.22337059 (Fig. 2, and S2) and at doi.org/10.6084/m9.figshare.28585775 (Fig. S5). Reads from the SWM1000 and parental strain are available on NCBI SRA under Bioproject PRJNA1235322

## Acknowledgements.

This research was supported by the U.S. Department of Agriculture National Institute of Food and Agriculture, Agriculture and Food Research Initiative (USDA-NIFA-AFRI Fellowship #: 2020-67012-31784 and Award #: 2023-67013-40245), Hatch Program (Project #1023861). A Research Experience for Post-Baccalaureate Students (REPS) supplement to out NSF Grant MCB 1716833 provided support for JKA. A portion of this research (JGI-SynBio 506458 Award DOI: 10.46936/10.25585/60000993) was performed under the JGI-EMSL Collaborative Science Initiative and used resources at the DOE Joint Genome Institute and the Environmental Molecular Sciences Laboratory, which are DOW Office of Science User Facilities. Both facilities are sponsored by the Office of Biological and Environmental Research and operated under Contract Nos. DE-AC02-05CH11231 (JGI) and DE-AC05-76RL01830 (EMSL).

---

**Fig. S1.**
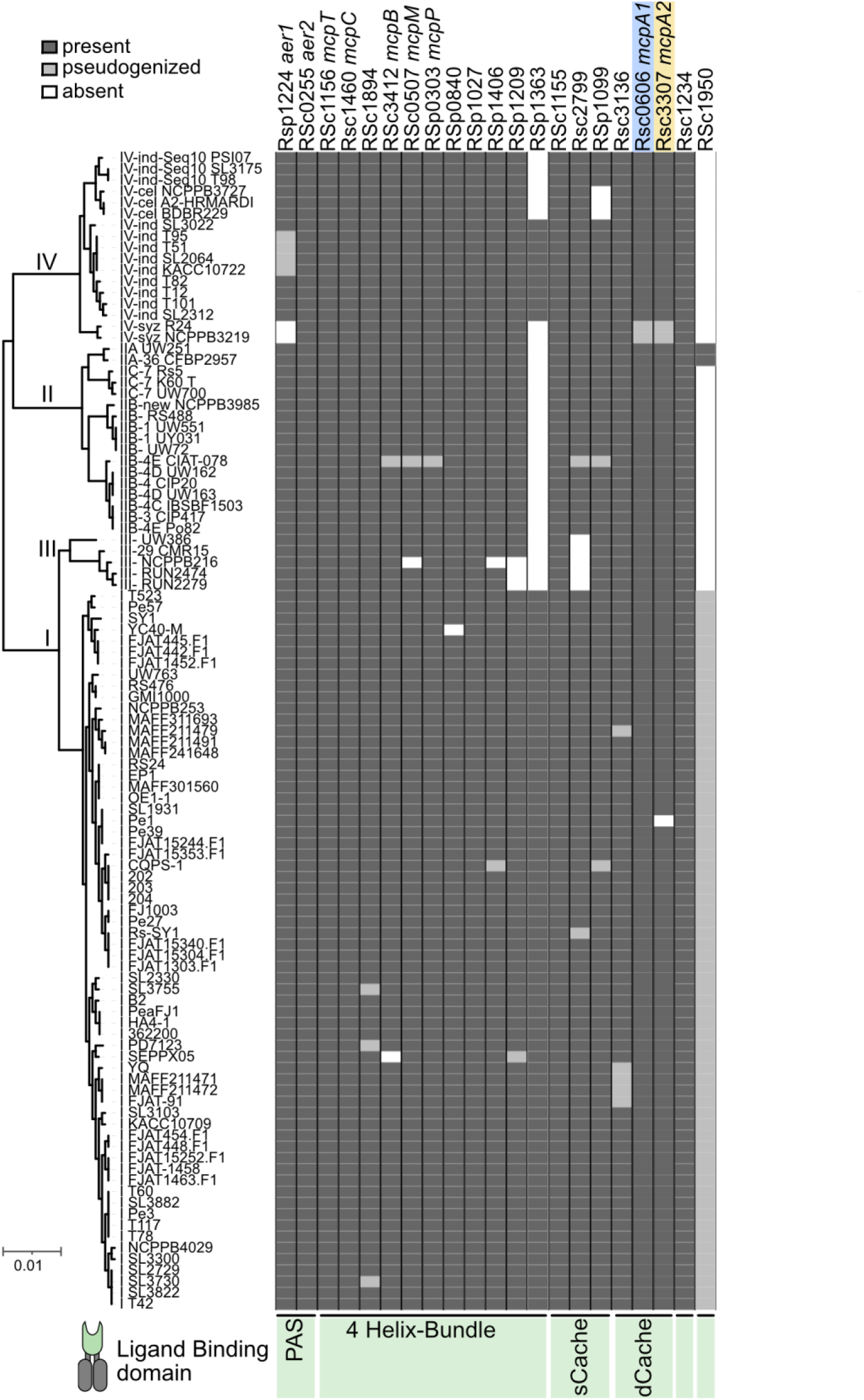
Detailed version of Figure 1 showing the names of strains and presence/absence of 21 *mcp* orthologs in the *Ralstonia* wilt pathogens.

**Fig. S2.**
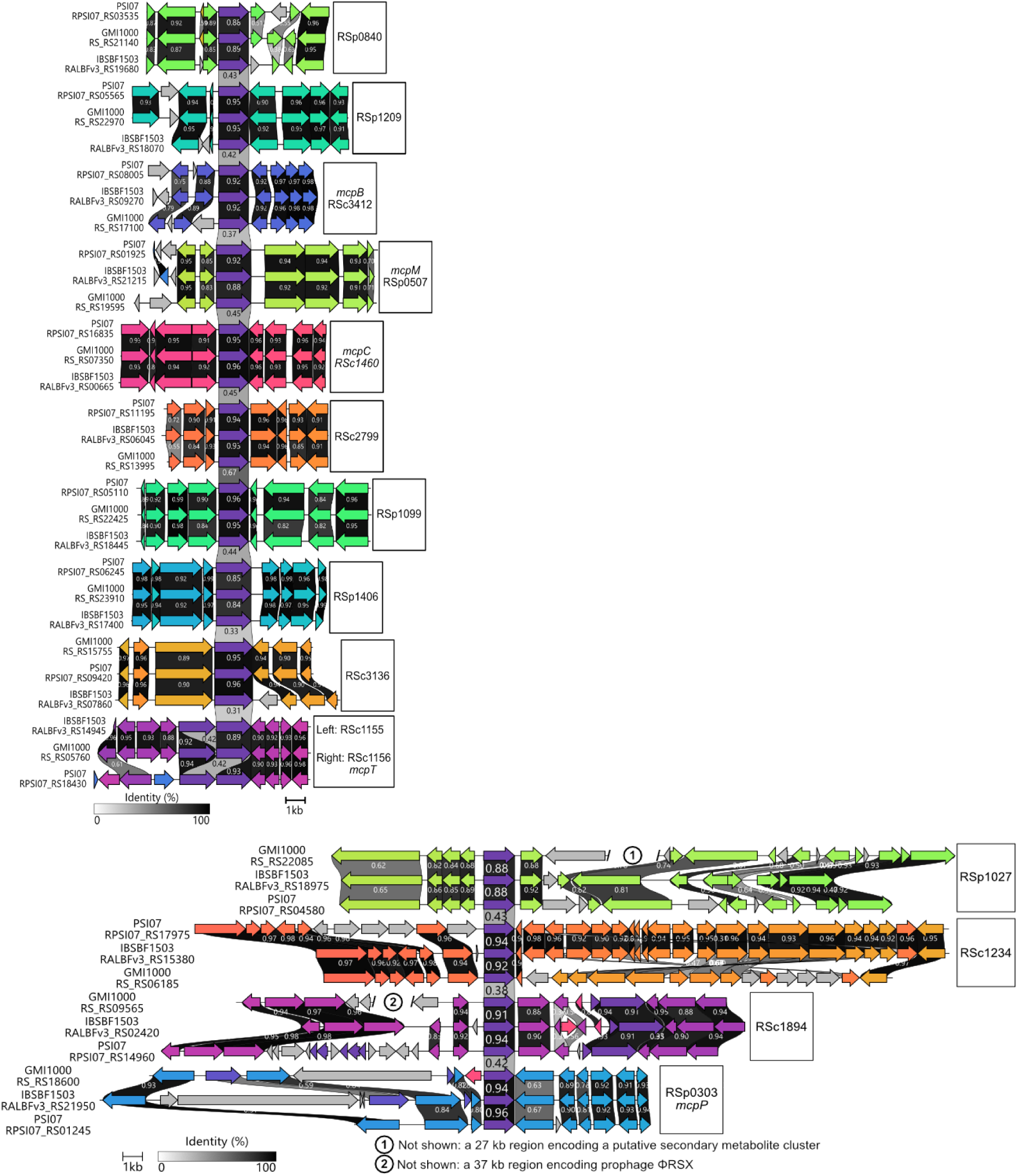
The genetic neighborhoods of 15 core *mcp* genes. The neighborhoods of *mcpA1* and *mcpA2* are shown in Fig. 2, and the neighborhoods for the energy taxis receptors were not visualized. Top: The gene neighborhoods of RSp0840, RSp1209, RSc3412 (*mcpB*), RSp0507 (*mcpM*), RSc1460 (*mcpC*), RSc2799, RSp1099, RSp1406, RSc3136, RSc1155, and RSc1156 (*mcpT*) are strongly conserved. Bottom: The genetic neighborhoods of RSp1027, RSc1234, RSc1894, and RSp0303 (*mcpP*) are generally conserved although there appear to be strain-specific gene insertions and inversions. Synteny was analyzed and visualized with Clinker (73). The interactive HTML files are available at doi.org/10.6084/m9.figshare.22337059

**Fig. S3.**
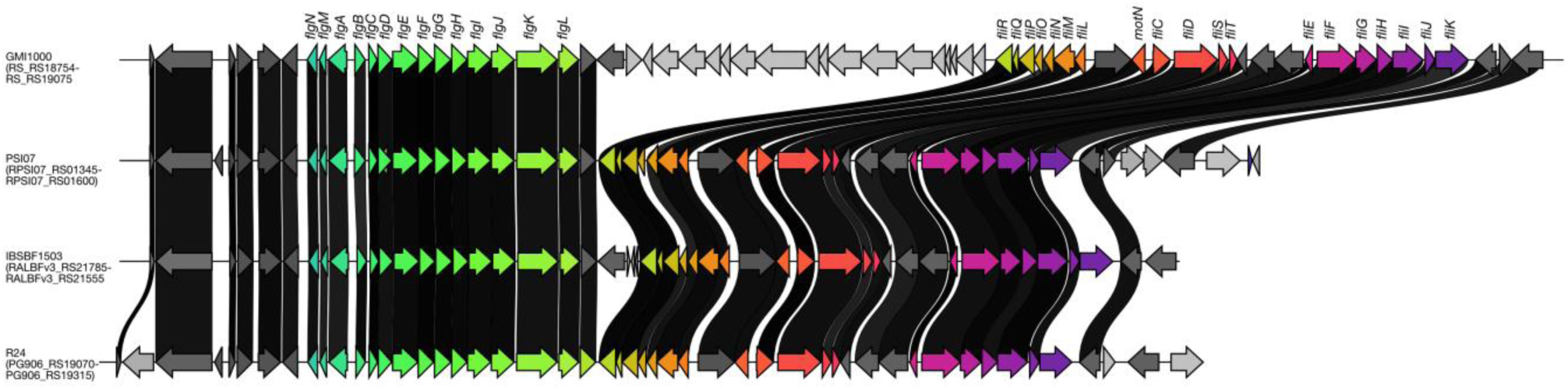
Synteny of *flgNMABCDEFGHIJKL, fliRQPONML, fliEFGHIJK,* and *motN* genes. Synteny is shown for *Ralstonia* model strains tested in this work (GMI1000, PSI07, and IBSBF1503) as well as SDC strain R24. Sequence identity greater than 90% is shown with black connecting bars.

**Fig. S4.**
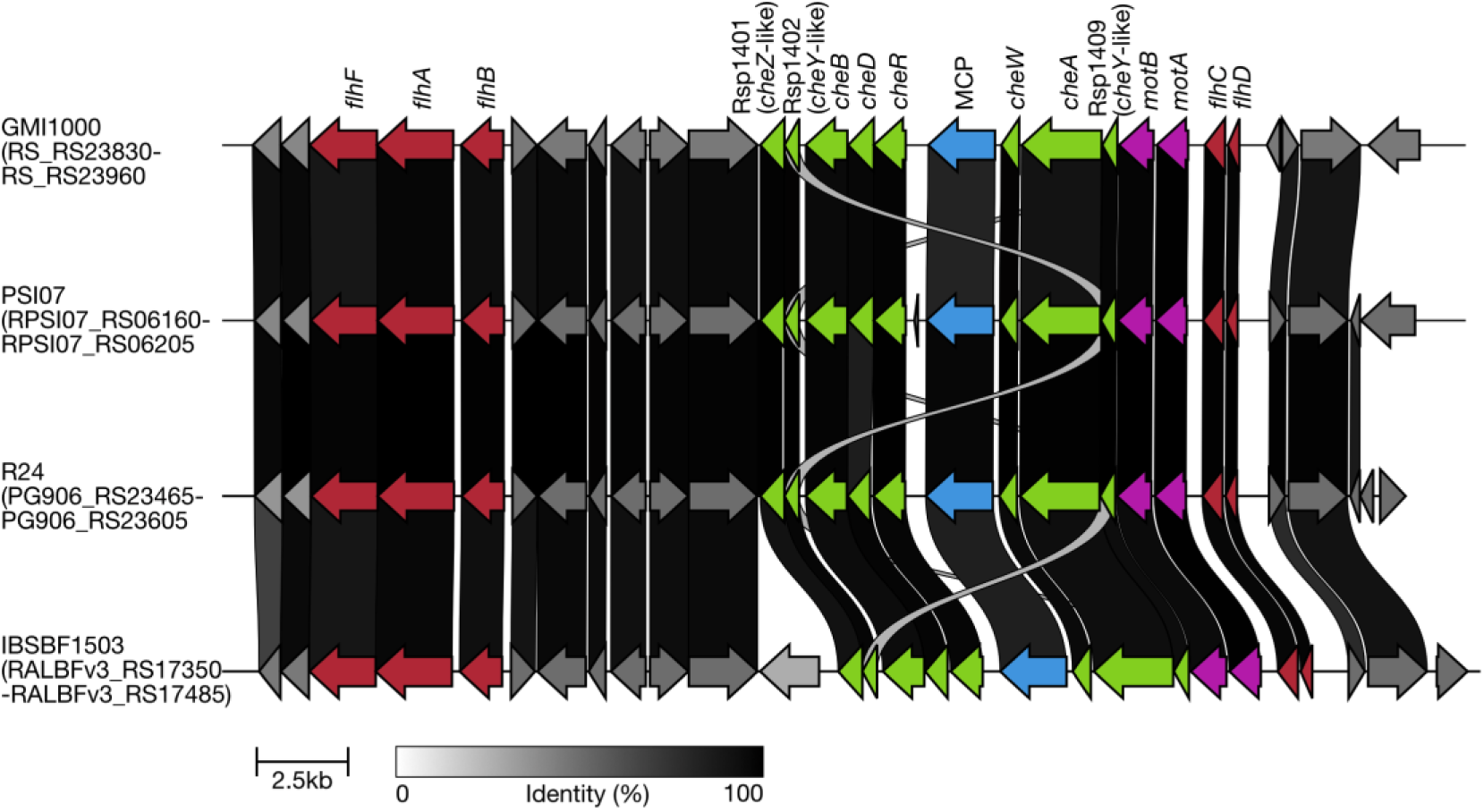
Genetic neighborhood encoding for the core chemotaxis machinery. This locus includes a *cheZ-* family gene, a *cheY*-family gene*, cheB, cheD, cheR, cheW,* a second *cheY*-family gene*, motA, motB*, *flhD, flhC, flhF, flhA,* and *flhB*. The *che* genes are highlighted in green, *mot* genes in fuchsia, *flh* genes in red, and the *mcp* gene RSp1406 in blue.

**Fig. S5.**
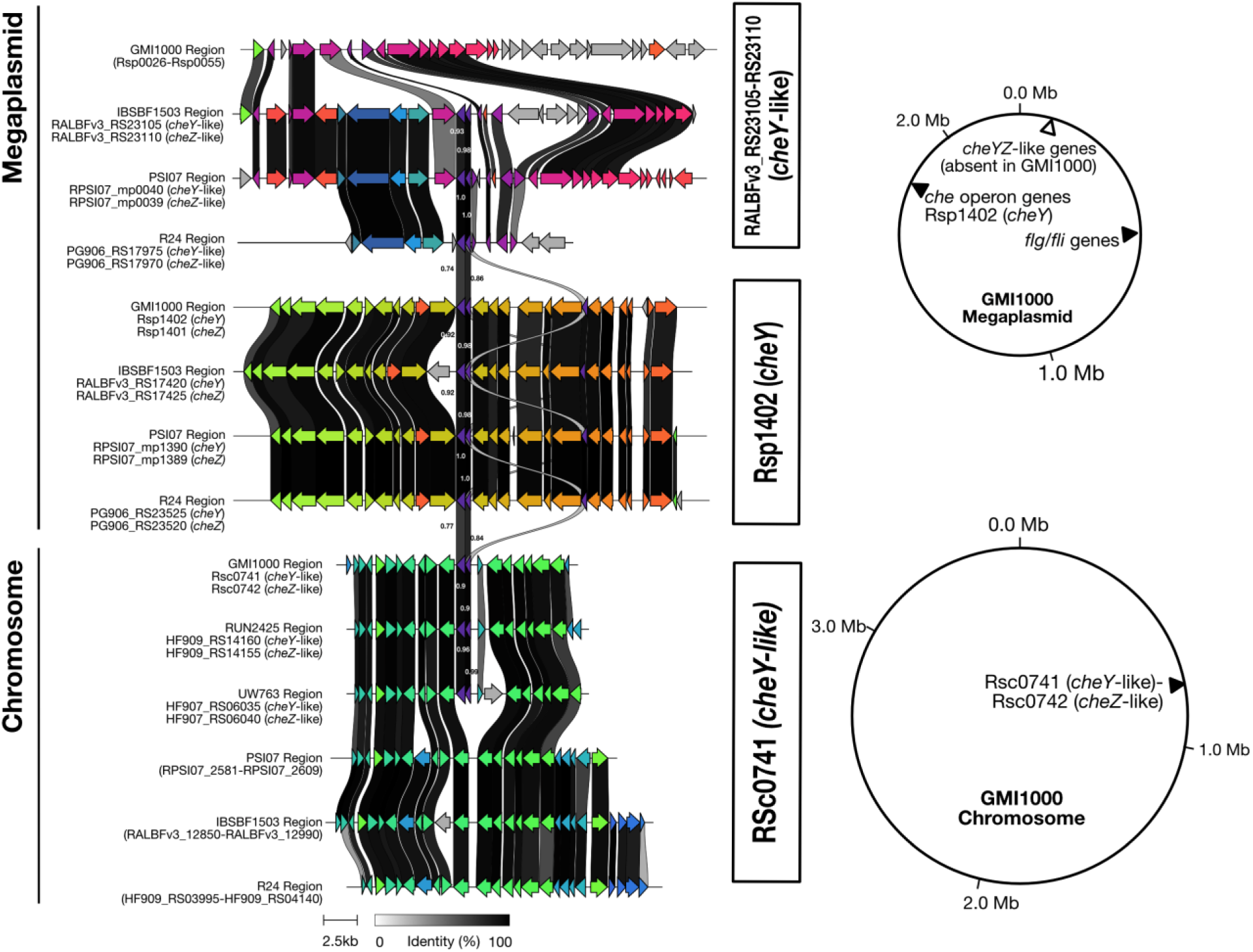
The genetic neighborhoods of *cheY*-like and *cheZ*-like genes. Synteny was analyzed for *cheY*-like genes in *Ralstonia* and visualized with Clinker (73). *cheYZ-*like genes are shown in purple in the synteny plots. *cheYZ (*RSp1401-Rsp1402), *cheY-* and *cheZ-*like gene clusters (RSc0741-RSc0742) are present in GMI1000 and examined phylotype I strains. A third set of *cheY-*like and *cheZ-*like genes (RALBFv3_RS23105 and RALBFv3_RS23110), are present in *R. solanacearum* and *R. syzygii*. The interactive HTML files for synteny analysis are available at doi.org/10.6084/m9.figshare.28585775.

**Fig. S6.**
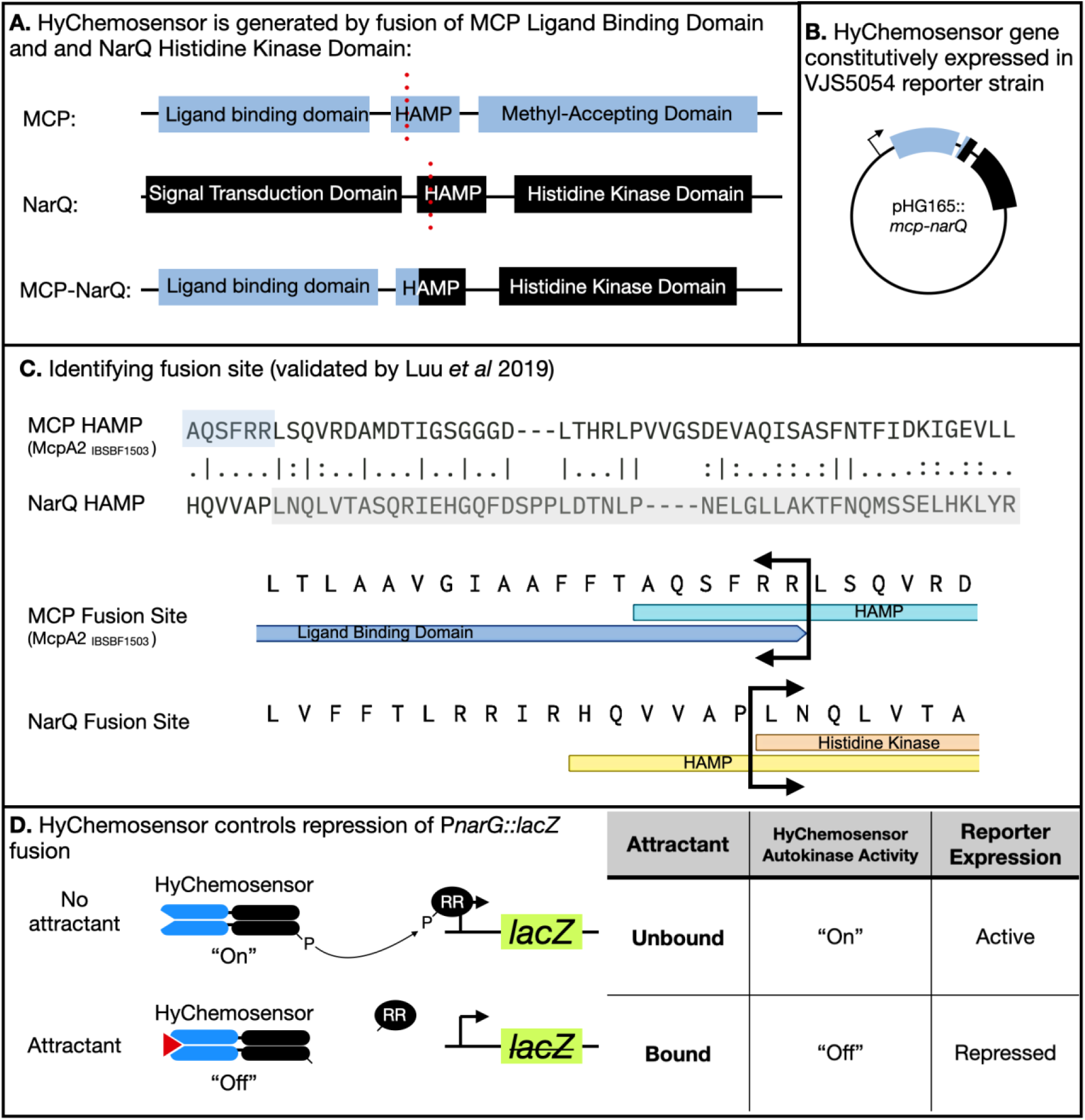
Construction and function of HyChemosensors. **A.** HyChemosensors are constructed by fusing region encoding the periplasmic ligand binding domain (PF12729) of an MCP gene with the cytoplasmic histidine kinase encoding domain (PF02518) of *E. coli narQ* at a conserved site in the HAMP domain (PF00672) that is present in both proteins. **B.** The HyChemosensor construct is constitutively expressed on the broad-host-range plasmid pHG165**. C.** Details of the HAMP fusion site for HyChemosensors. The protein sequences of the HAMP domains of McpA2 (IBSBF1503) and NarQ are shown. Top: Alignment of the HAMP domain regions of McpA2 and NarQ. Bottom: Cartoon schematics of the primary sequence and domains of McpA2 and NarQ. The translational fusion site is labeled in all panels. **D.** The HyChemosensor protein controls the repression of a genomic *PnarG*-*lacZ* transcriptional fusion. When the HyChemosensor is not bound to an attractant, the histidine kinase domain is auto-phosphorylated in an “On” state. The phosphorylation signal is transduced to the response regulator (RR) NarL. Phosphorylated NarL binds the operator of *PnarG* activating expression of *lacZ*. When an attractant binds to the HyChemosensor, the dephosphorylated NarL will not bind to the operator of *PnarG* and *lacZ* expression not activated. Thus, *lacZ* is expressed when there is no attractant, and the presence of an attractant reduces *lacZ* expression.

**Fig. S7.**
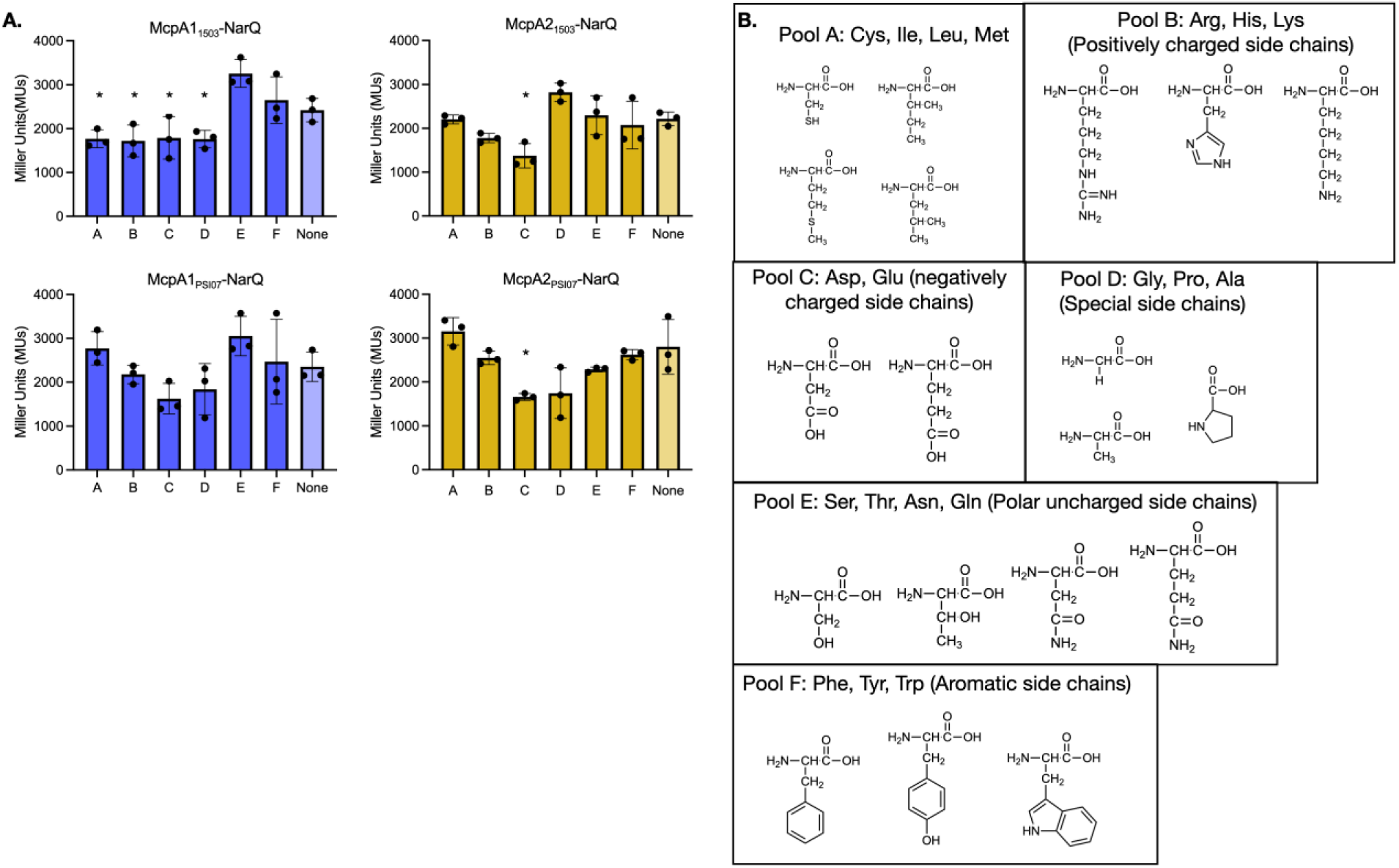
**A.** Responses of McpA1-NarQ (blue) and McpA2-NarQ (yellow) orthologs to various pools of amino acids are shown. Composition of amino acid pools A-F and their structural motifs are shown in **B.** The final concentration for each amino acid within the pool was 1 mM. Valine was excluded because it arrested growth of *E. coli* VJS5054 in MSB. Experimental averages of 3 independent experiments are shown and standard deviations are represented with error bars.

**Fig. S8.**
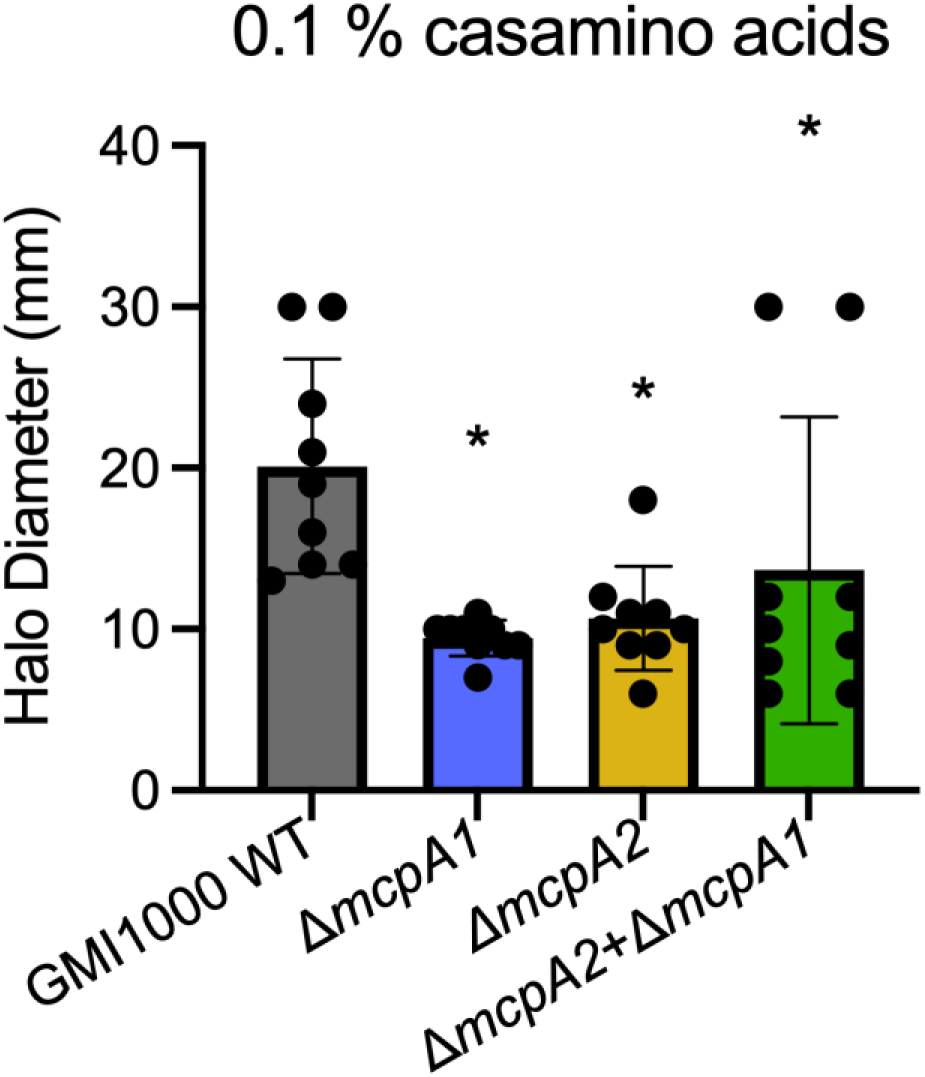
McpA1 and McpA2 contribute to *Ralstonia* chemotaxis in rich media soft agar. WT GMI1000, Δ*mcpA1* mutant, Δ*mcpA2* mutant, or Δ*mcpA1* Δ*mcpA2* double mutant strains were inoculated into soft agar plates containing 0.1% w/v casamino acids. The diameters of the halos were measured after incubation at 28°C for 50-60 hr. Data points show two trials with 4-5 biological replicates per trial. The asterisks indicate significance, relative to the WT control by the Krusker-Wallis test, (p<0.05).

**Fig. S9.**
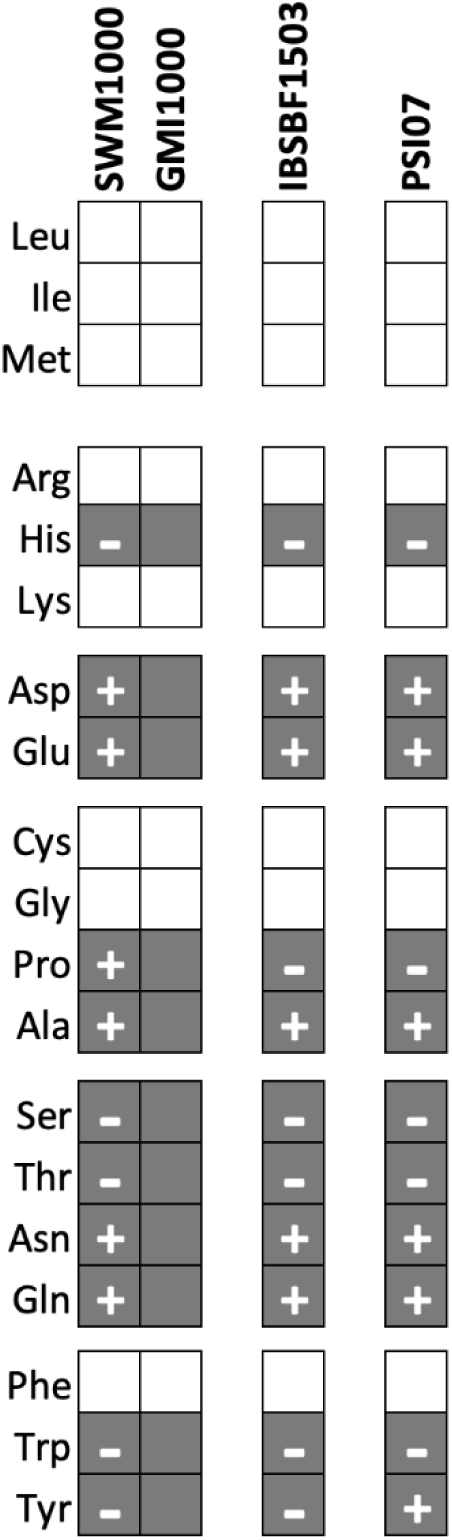
Growth and attraction to proteinogenic amino acids as sole carbon sources by SWM1000, GMI1000, IBSBF1503, and PSI07. A white box indicates no growth was observed after 72 hr at 28° C in minimal medium supplemented with 1 mM of the tested amino acid. Gray boxes indicate growth was observed. For amino acids that could be utilized as a carbon source, a plus (+) indicates that this amino acid also elicited a chemoattractive response to the amino acid in soft agar swim plate assays, whereas a minus (-) indicates that no chemoattractive response to these amino acids was observed after 84 hr incubation. GMI1000 was not tested for amino acid responses because the response time was too long. Gray boxes without a (+) or (-) indicate that growth was observed, but chemotaxis was not reliably testable by this assay. Three separate colonies of each strain were tested in three replicate experiments.

**Fig. S10.**
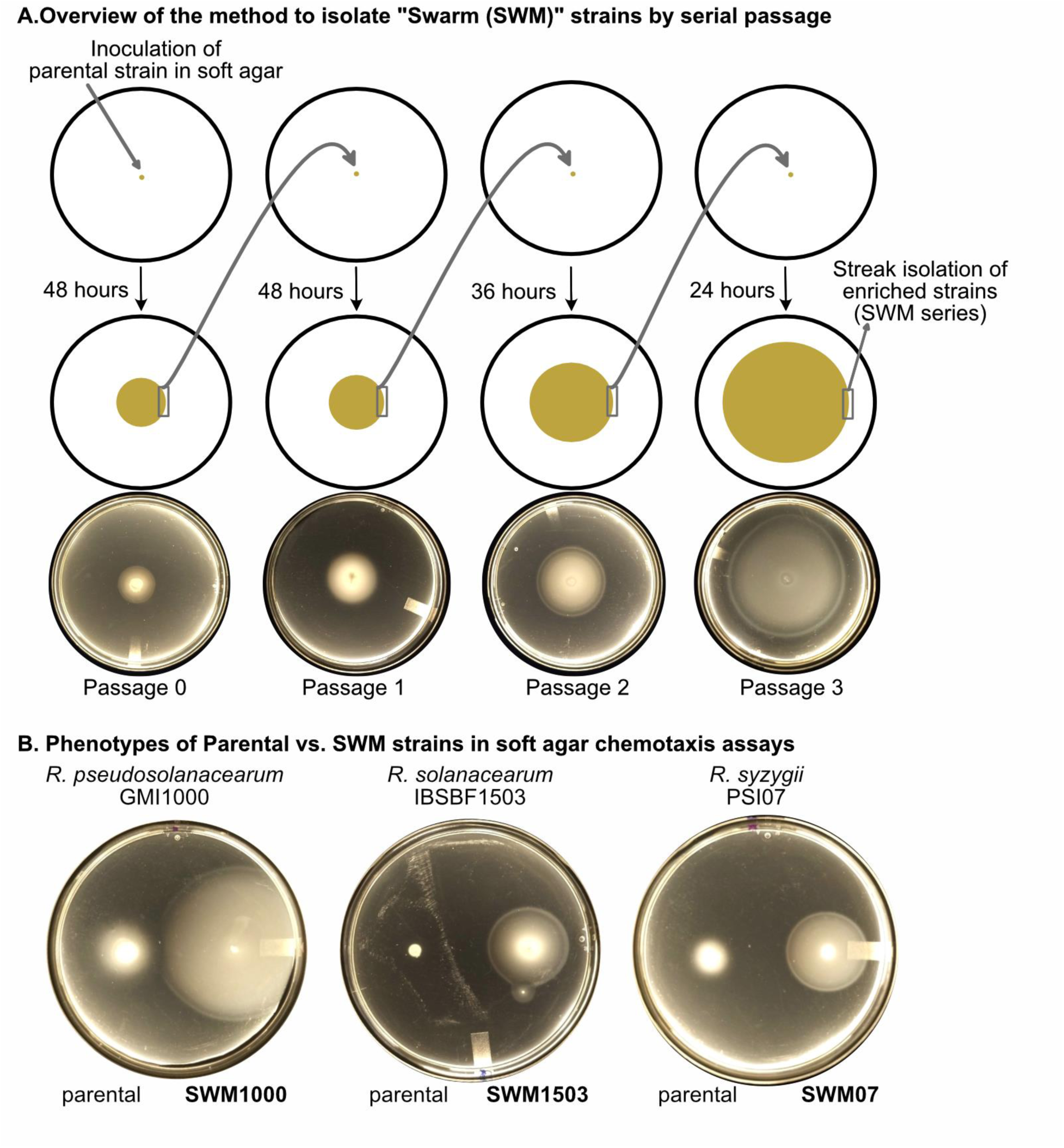
A. Schematic cartoon (top) and representative images (bottom) of the serial passage approach to enrich *Ralstonia* Swarm (SWM) strains for an enhanced chemotactic response. Photos show the isolation of the IBSBF1503 derivative SWM1503 (B) Representative qualitative swim plates showing enhanced chemotactic response of SWM strains over their parental strain. Each strain was resuscitated from cryostorage and then inoculated into fresh 50% CPG containing 0.25% agar to confirm genetic stability of the selected phenotype.

**Fig S11.**
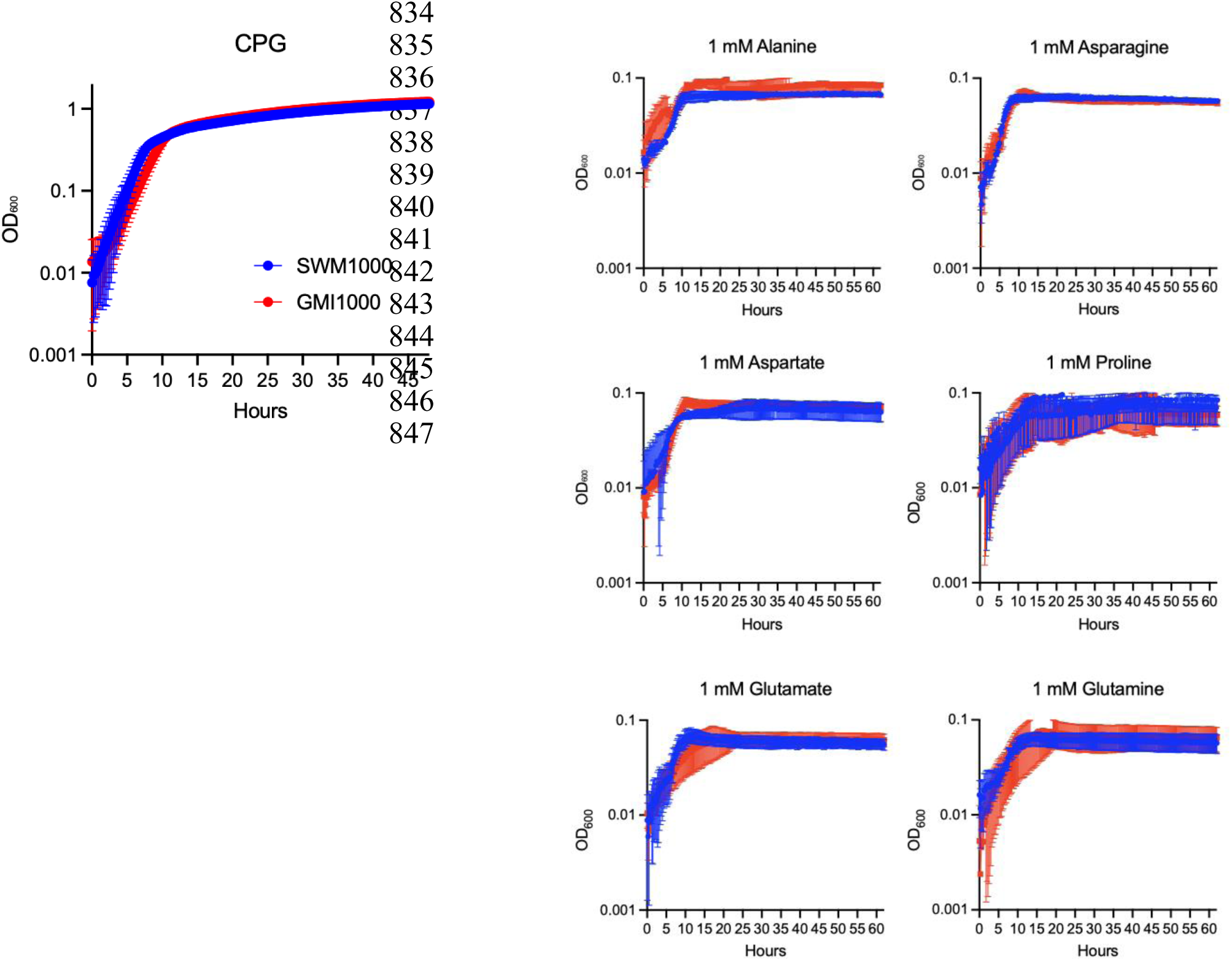
Growth of SWM1000 (blue) and GMI1000 (red) in rich medium (CPG) and minimal medium (MSB) with the indicated amino acids as sole carbon sources. Data points represent the averages of three individually cultivated colonies. Error bars represent standard deviations.

**Fig. S12.**
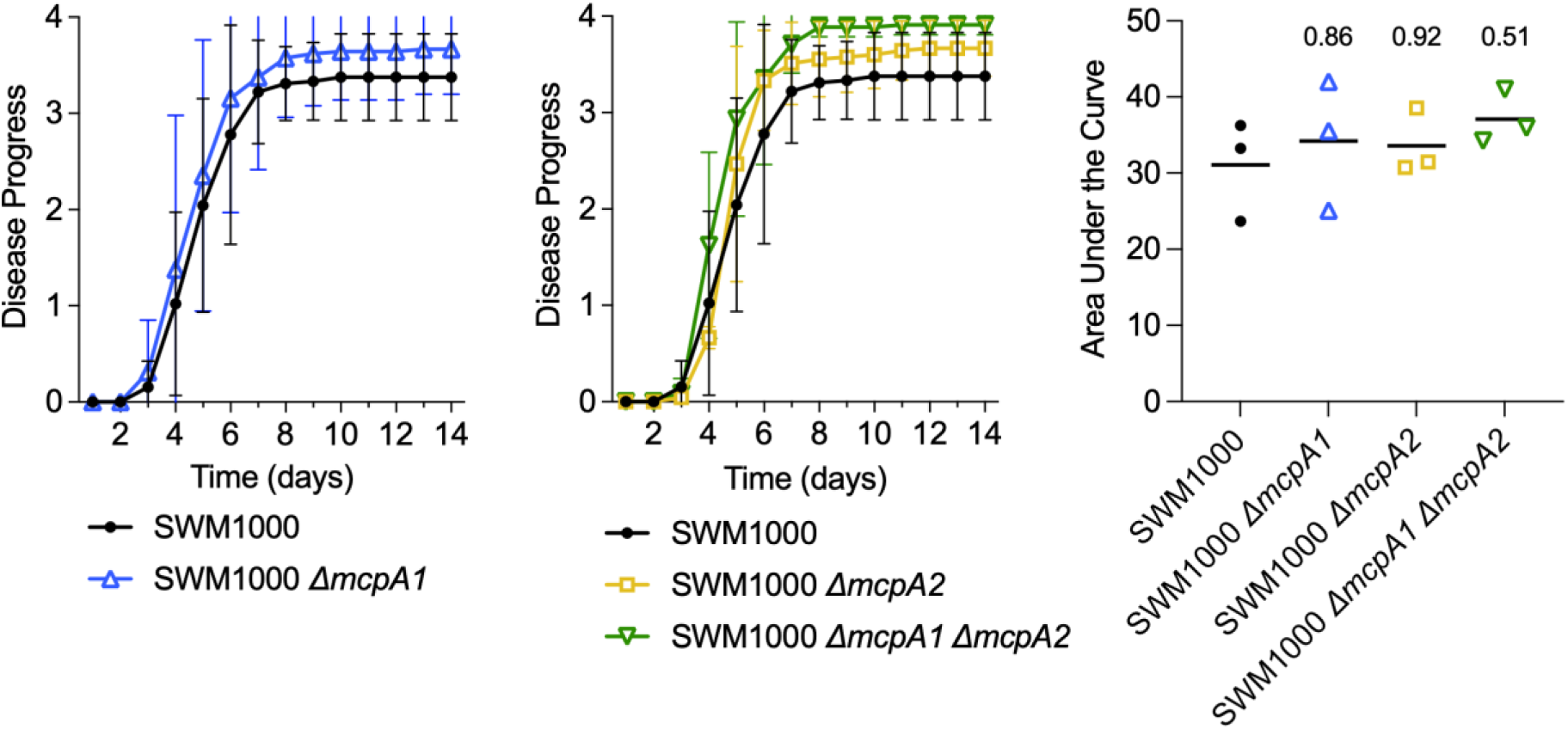
McpA2 does not contribute to *Ralstonia* virulence after cut petiole inoculation. Tomato cv. moneymaker plants were directly inoculated via cut petioles with the swarmed strains SWM1000, Δ*mcpA1*, Δ*mcpA2*, or Δ*mcpA1* Δ*mcpA2*. Plants were rated daily on a disease index scale from 0 to 4 for 14 days. Each data point is the average disease index for that day across three independent trials (n=15 plants per trial). These data are also represented as an area under the curve for each replicate trail. P-values shown represent the output of Ordinary One-Way ANOVA statistical tests.

**Figure S13.**
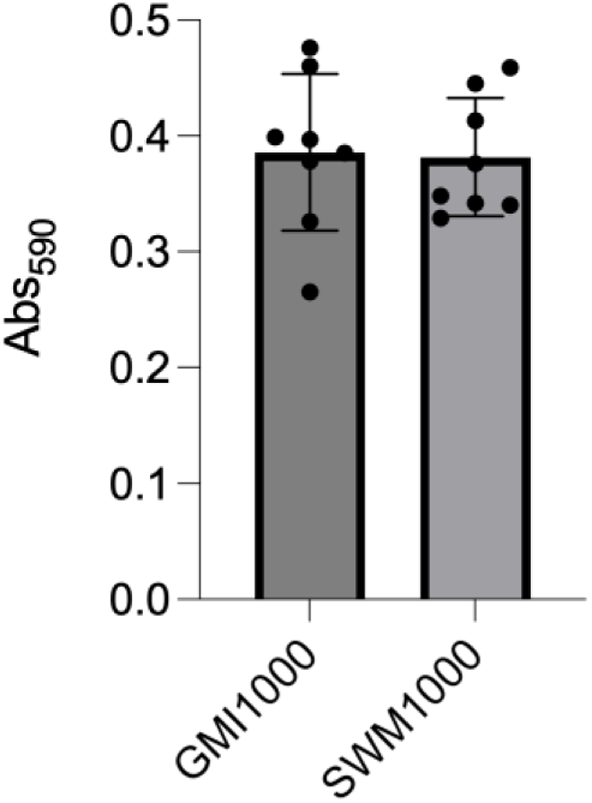
Quantification of biofilm accumulation on PVC of GMI1000 and SWM1000 after culturing in CPG medium. Bars represent the averages of eight individually cultivated colonies. Error bars represent standard deviation of the data. A student’s t-test indicated no statistical difference between the two strains.

**Table S1.** Metadata describing the MCPs in the *Ralstonia* wilt pathogens.

| Gene name: | GMI1000 locus (original): | GMI1000 locus (NCBI): | Core: | Ligand binding domain: | Contains AA-sensing domain ("YxxxxRxWY(~8x)Y(~25x)D"): | Chemicals sensed: | Citation |
| --- | --- | --- | --- | --- | --- | --- | --- |
| *mcpA1* | RSc0606 | RS_RS03035 | Yes | double cache | Yes | amino acids (+) | (16) and in this report |
| *mcpA2* | RSc3307 | RS_RS16570 | Yes | double cache | Yes | amino acids (+) | This report |
| *mcpB* | RSc3412 | RS_RS17100 | Yes | 4-helix bundle (4HB) | No | Boric Acid (+) | (60) |
| *mcpC* | RSc1460 | RS_RS07350 | Yes | 4-helix bundle (4HB) | No | Citrate (+) | (18) |
| *mcpM* | RSp0507 | RS_RS19595 | Yes | 4-helix bundle (4HB) | No | L-malate (+), D-malate (+) | (16, 17) |
| *mcpP* | RSp0303 | RS_RS18600 | Yes | 4-helix bundle (4HB) | No | Citrate (+), phosphate (+), maleate (-) | (18) |
| *mcpT* | RSc1156 | RS_RS05760 | Yes | 4-helix bundle (4HB) | No | L-tartrate (+), D-malate (+) | (17) |
| RSc1155 | RSc1155 | RS_RS05755 | Yes | single cache | No | unknown |  |
| RSc1234 | RSc1234 | RS_RS06185 | Yes | unclassified domain | No | unknown |  |
| RSc1894 | RSc1894 | RS_RS09565 | Yes | 4-helix bundle (4HB) | No | unknown |  |
| RSc1950 | RSc1950 | RS_RS09805 | no | unclassified domain | No | unknown |  |
| RSc2799 | RSc2799 | RS_RS13995 | Yes | single cache | No | unknown |  |
| RSc3136 | RSc3136 | RS_RS15755 | Yes | double cache | No | unknown |  |
| RSp0840 | RSp0840 | RS_RS21140 | Yes | 4-helix bundle (4HB) | No | unknown |  |
| RSp1027 | RSp1027 | RS_RS22085 | Yes | 4-helix bundle (4HB) | No | unknown |  |
| RSp1099 | RSp1099 | RS_RS22425 | Yes | single cache | No | unknown |  |
| RSp1209 | RSp1209 | RS_RS22970 | Yes | 4-helix<br>bundle<br>(4HB) | No | unknown |  |
| RSp1363 | RSp1363 | RS_RS23695 | no | 4-helix<br>bundle<br>(4HB) | No | unknown |  |
| RSp1406 | RSp1406 | RS_RS23910 | Yes | 4-helix<br>bundle<br>(4HB) | No | unknown |  |
| *aer1* | RSp1224 | RS_RS23045 | Yes | PAS | No | energy taxis | (3) |
| *aer2* | RSp0255 | RS_RS18385 | Yes | PAS | No | energy taxis | (3) |

**Table S2.** Strains used in this study.

| Strain: | Background: | Genotype: | Reference: |
| --- | --- | --- | --- |
| GMI1000 | *Ralstonia pseudosolanacearum* isolate | Wildtype isolate | (78) |
| IBSBF1503 | *Ralstonia solanacearum* isolate | Wildtype isolate | (79, 80) |
| PSI07 | *Ralstonia syzygii* isolate | Wildtype isolate | (79, 80) |
| SWM1000 | Swarm-enriched *Ralstonia pseudosolanacearum* GMI1000 | Swm+ enriched isolate | This report |
| SWM1503 | Swarm-enriched *Ralstonia solanacearum* IBSBF1503 | Swm+ enriched isolate | This report |
| SWM07 | Swarm-enriched *Ralstonia syzygii* PSI07 | Swm+ enriched isolate | This report |
| JKA004 | GMI1000 | $\Delta mcpA1$ (RSc0606) | This report |
| JKA005 | GMI1000 | $\Delta mcpA2$ (RSc3307) | This report |
| JKA007 | GMI1000 | $\Delta mcpA1$ (RSc0606), $\Delta mcpA2$ (RSc3307) | This report |
| JKA008 | SWM1000 | Swm+, $\Delta mcpA2$ (RSc3307) | This report |
| JKA009 | SWM1000 | Swm+, $\Delta mcpA1$ (RSc0606) | This report |
| JKA010 | SWM1000 | Swm+, $\Delta mcpA1$ (RSc0606), $\Delta mcpA2$ (RSc3307) | This report |
| VJS5054 | *E. coli* | $\lambda\Phi(narG-lacZ)250 \Delta(argF-lac)U169 \Delta narX242 narQ251::Tn10d(Tc) pcnB1 zad-981::Tn10d(Km)$ | (30) |
| DH5α | *E. coli* | Cloning strain | Life Technologies, Gaithersburg, MD |

**Table S3.** Plasmids used in this study.

| Name: | Description: | Purpose: | Reference: |
| --- | --- | --- | --- |
| pVJS3354 | *narQ* in pHG165, Amp<sup>R</sup> | Cloning and expression plasmid containing full-length *narQ* | (28) |
| pBSR8 | pVJS3354::*mcpA2*<sub>PSI07</sub>-*narQ*<br>LBD-coding sequence of *mcpA2* originating from PSI07 with cytoplasmic and HAMP domain-coding sequences of *narQ* in pHG165, Amp<sup>R</sup> | MpcA2<sub>PSI07</sub>-NarQ HyChemosensor; Testing ligand-binding of McpA2 | This report |
| pBSR16 | pVJS3354::*mcpA2*<sub>IBSBF1503</sub>-*narQ*<br>LBD-coding sequence of *mcpA2* originating from IBSBF1503 with cytoplasmic and HAMP domain-coding sequences of *narQ* in pHG165, Amp<sup>R</sup> | MpcA2<sub>1503</sub>-NarQ HyChemosensor; Testing ligand-binding of McpA2 | This report |
| pBSR21 | pVJS3354::*mcpA1*<sub>IBSBF1503</sub>-*narQ*<br>LBD-coding sequence of *mcpA1* originating from IBSBF1503 fused with cytoplasmic and HAMP domain-coding sequences of *narQ* in pHG165, Amp<sup>R</sup> | McpA1<sub>1503</sub>-NarQ HyChemosensor; Testing ligand-binding of McpA1 | This report |
| pBSR1 | pVJS3354::*mcpA1*<sub>PSI07</sub>-*narQ*<br>LBD-coding sequence of *mcpA1* originating from PSI07 fused with cytoplasmic and HAMP domain-coding sequences of *narQ* in pHG165, Amp<sup>R</sup> | MpcA1<sub>PSI07</sub>-NarQ HyChemosensor; Testing ligand-binding of McpA1 | This report |
| pJPF017 | LBD-coding sequence of *mcfA* fused with cytoplasmic and HAMP domain-coding sequences of *narQ* in pHG165, Amp<sup>R</sup> | McfA-NarQ HyChemosensor; Control for MCP-NarQ fusions | (25) |
| pUFR80 | *sucS* (*sacB*), Km<sup>R</sup> | Markerless deletion via selection/counterselection | (81) |
| pUFR80::3307_delGMI1000 | *RSc3307* (*mcpA2*) upstream and downstream 1-kb PCR fragments fused and cloned pUFR80; Km<sup>R</sup> | to delete *mcpA2* (*RSc3307*) | This report |
| pUFR80::mcpA1_delGMI1000 | *RSc0606* (*mcpA2*) upstream and downstream 1-kb PCR fragments fused and cloned pUFR80; Km<sup>R</sup> | to delete *mcpA1* (*RSc0606*) | This report |

**Table S4.**
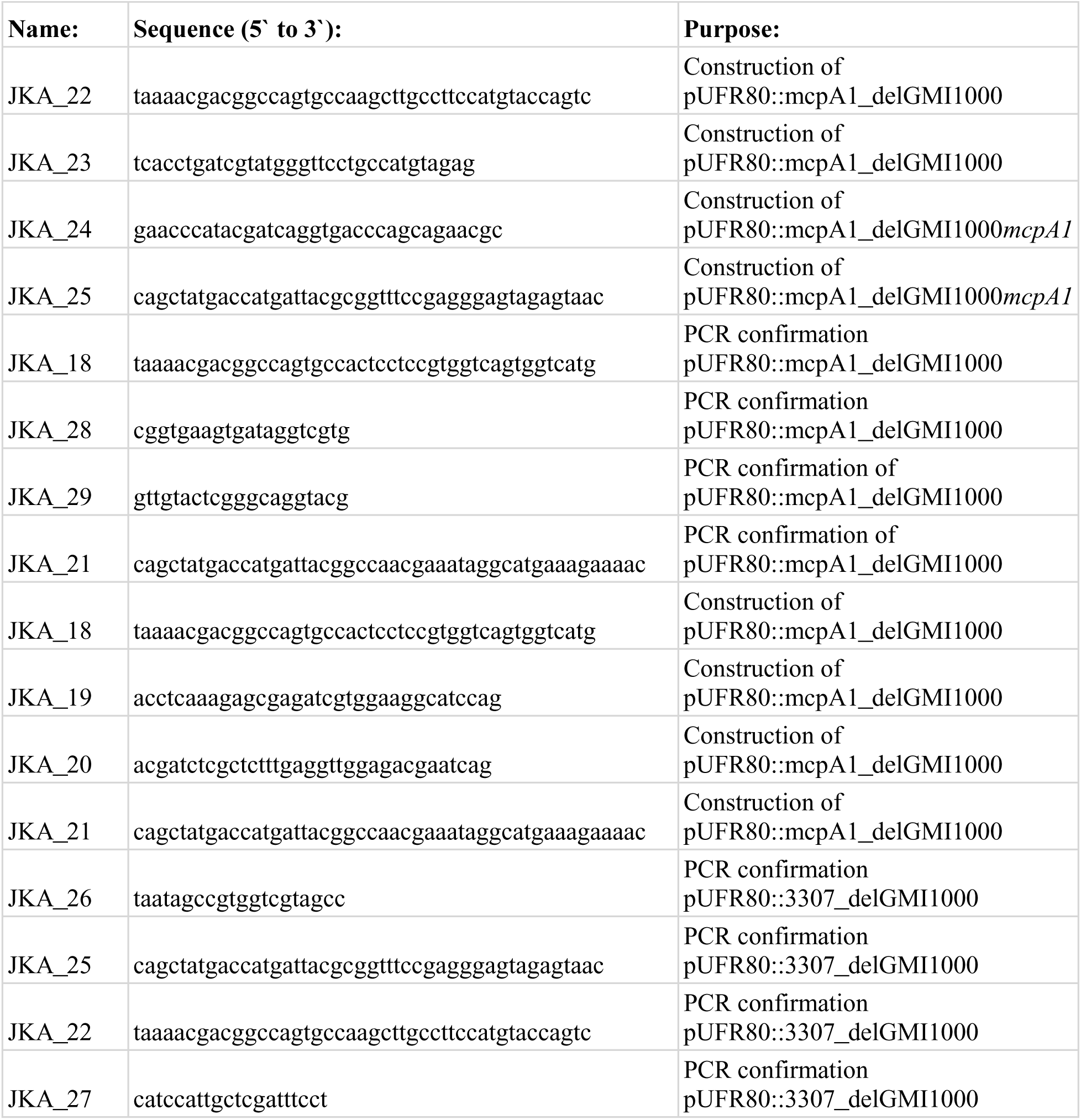
Primers used in this study.

## References

1. Lowe-Power T, Avalos J, Munoz MC, Chipman K, Williams D. 2020. A meta-analysis of the known global distribution and host range of the Ralstonia species complex.

2. Vives-Peris V, de Ollas C, Gómez-Cadenas A, Pérez-Clemente RM. 2020. Root exudates: from plant to rhizosphere and beyond. Plant Cell Rep 39:3–17.

3. Yao J, Allen C. 2007. The plant pathogen *Ralstonia solanacearum* needs aerotaxis for normal biofilm formation and interactions with its tomato host. J Bacteriol 189:6415–6424.

4. Yao J, Allen C. 2006. Chemotaxis is required for virulence and competitive fitness of the bacterial wilt pathogen *Ralstonia solanacearum*. J Bacteriol 188:3697–3708.

5. Jacob-Dubuisson F, Mechaly A, Betton J-M, Antoine R. 2018. Structural insights into the signalling mechanisms of two-component systems. Nat Rev Microbiol 16:585–593.

6. Bi S, Sourjik V. 2018. Stimulus sensing and signal processing in bacterial chemotaxis. Curr Opin Microbiol 45:22–29.

7. Sourjik V, Wingreen NS. 2012. Responding to chemical gradients: bacterial chemotaxis. Curr Opin Cell Biol 24:262–268.

8. Gegner JA, Graham DR, Roth AF, Dahlquist FW. 1992. Assembly of an MCP receptor, CheW, and kinase CheA complex in the bacterial chemotaxis signal transduction pathway. Cell 70:975–982.

9. Wadhams GH, Armitage JP. 2004. Making sense of it all: Bacterial chemotaxis. Nat Rev Mol Cell Biol 5:1024–1037.

10. Sampedro I, Parales RE, Krell T, Hill JE. 2015. *Pseudomonas* chemotaxis. FEMS Microbiol Rev 39:17–46.

11. Ortega Á, Zhulin IB, Krell T. 2017. Sensory repertoire of bacterial chemoreceptors. Microbiol Mol Biol Rev 81:1–28.

12. Sanchis-López C, Cerna-Vargas JP, Santamaría-Hernando S, Ramos C, Krell T, Rodríguez-Palenzuela P, López-Solanilla E, Huerta-Cepas J, Rodríguez-Herva JJ. 2021. Prevalence and specificity of chemoreceptor profiles in plant-associated bacteria. mSystems 6:e0095121.

13. Tans-Kersten J, Huang H, Allen C. 2001. *Ralstonia solanacearum* needs motility for invasive virulence on tomato. J Bacteriol 183:3597–3605.

14. Lacal J, García-Fontana C, Muñoz-Martínez F, Ramos JL, Krell T. 2010. Sensing of environmental signals: Classification of chemoreceptors according to the size of their ligand binding regions. Environ Microbiol 12:2873–2884.

15. Matilla MA, Krell T. 2024. Sensing the environment by bacterial plant pathogens: What do their numerous chemoreceptors recognize? Microb Biotechnol 17:e14368.

16. Hida A, Oku S, Kawasaki T, Nakashimada Y, Tajima T, Kato J. 2015. Identification of the *mcpA* and *mcpM* genes, encoding methyl-accepting proteins involved in amino acid and l-malate chemotaxis, and involvement of McpM-mediated chemotaxis in plant infection by *Ralstonia pseudosolanacearum* (formerly *Ralstonia solanacearum* phylotypes I and III). Appl Environ Microbiol 81:7420–7430.

17. Tunchai M, Hida A, Oku S, Nakashimada Y, Tajima T, Kato J. 2017. Identification and characterization of chemosensors for d-malate, unnatural enantiomer of malate, in *Ralstonia pseudosolanacearum*. Microbiology 163:233–242.

18. Hida A, Tajima T, Kato J. 2019. Two citrate chemoreceptors involved in chemotaxis to citrate and/or citrate-metal complexes in *Ralstonia pseudosolanacearum*. J Biosci Bioeng 127:169–175.

19. Hughes JG, Zhang X, Parales JV, Ditty JL, Parales RE. 2017. *Pseudomonas putida* F1 uses energy taxis to sense hydroxycinnamic acids. Microbiology 163:1490–1501.

20. Rabinovitch-Deere CA, Parales RE. 2012. Three types of taxis used in the response of *Acidovorax* sp. strain JS42 to 2-nitrotoluene. Appl Environ Microbiol 78:2306–2315.

21. Schweinitzer T, Josenhans C. 2010. Bacterial energy taxis: a global strategy? Arch Microbiol 192:507–520.

22. Velando F, Matilla MA, Zhulin IB, Krell T. 2023. Three unrelated chemoreceptors provide *Pectobacterium atrosepticum* with a broad-spectrum amino acid sensing capability. Microb Biotechnol 16:1548–1560.

23. Parales RE, Luu RA, Chen GY, Liu X, Wu V, Lin P, Hughes JG, Nesteryuk V, Parales JV, Ditty JL. 2013. *Pseudomonas putida* F1 has multiple chemoreceptors with overlapping specificity for organic acids. Microbiology 159:1086–1096.

24. Schwarzer C, Fischer H, Machen TE. 2016. Chemotaxis and binding of *Pseudomonas aeruginosa* to scratch-wounded human cystic fibrosis airway epithelial cells. PLoS One 11:e0150109.

25. Luu RA, Schomer RA, Brunton CN, Truong R, Ta AP, Tan WA, Parales JV, Wang Y-J, Huo Y-W, Liu S-J, Ditty JL, Stewart V, Parales RE. 2019. Hybrid Two-Component Sensors for Identification of Bacterial Chemoreceptor Function. Appl Environ Microbiol 85.

26. Gumerov VM, Andrianova EP, Matilla MA, Page KM, Monteagudo-Cascales E, Dolphin AC, Krell T, Zhulin IB. 2022. Amino acid sensor conserved from bacteria to humans. Proc Natl Acad Sci U S A 119:e2110415119.

27. Corral-Lugo A, De la Torre J, Matilla MA, Fernández M, Morel B, Espinosa-Urgel M, Krell T. 2016. Assessment of the contribution of chemoreceptor-based signalling to biofilm formation. Environ Microbiol 18:3355–3372.

28. Appleman JA, Chen LL, Stewart V. 2003. Probing conservation of HAMP linker structure and signal transduction mechanism through analysis of hybrid sensor kinases. J Bacteriol 185:4872– 4882.

29. Stewart V. 1993. Nitrate regulation of anaerobic respiratory gene expression in *Escherichia coli*. Mol Microbiol 9:425–434.

30. Williams SB, Stewart V. 1997. Nitrate- and nitrite-sensing protein NarX of *Escherichia coli* K-12: mutational analysis of the amino-terminal tail and first transmembrane segment. J Bacteriol 179:721–729.

31. Ditty JL, Parales RE. 2017. Protocols for the Measurement of Bacterial Chemotaxis to Hydrocarbons, p. 7–42. In McGenity, TJ, Timmis, KN, Nogales, B (eds.), Hydrocarbon and Lipid Microbiology Protocols: Activities and Phenotypes. Springer Berlin Heidelberg, Berlin, Heidelberg.

32. Harwood CS, Rivelli M, Ornston LN. 1984. Aromatic acids are chemoattractants for *Pseudomonas putida*. J Bacteriol 160:622–628.

33. Harwood CS, Nichols NN, Kim MK, Ditty JL, Parales RE. 1994. Identification of the pcaRKF gene cluster from *Pseudomonas putida*: involvement in chemotaxis, biodegradation, and transport of 4-hydroxybenzoate. J Bacteriol 176:6479–6488.

34. Huang Z, Wang Y-H, Zhu H-Z, Andrianova EP, Jiang C-Y, Li D, Ma L, Feng J, Liu Z-P, Xiang H, Zhulin IB, Liu S-J. 2019. Cross talk between chemosensory pathways that modulate chemotaxis and biofilm formation. MBio 10.

35. Rossi E, Paroni M, Landini P. 2018. Biofilm and motility in response to environmental and host-related signals in Gram negative opportunistic pathogens. J Appl Microbiol 125:1587–1602.

36. Meng F, Yao J, Allen C. 2011. A *motN* mutant of *Ralstonia solanacearum* is hypermotile and has reduced virulence. J Bacteriol 193:2477–2486.

37. Khokhani D, Lowe-Power TM, Tran TM, Allen C. 2017. A Single Regulator Mediates Strategic Switching between Attachment/Spread and Growth/Virulence in the Plant Pathogen *Ralstonia solanacearum*. MBio 8.

38. Deatherage DE, Barrick JE. 2014. Identification of mutations in laboratory-evolved microbes from next-generation sequencing data using breseq. Methods Mol Biol 1151:165–188.

39. Lynch JM, Whipps JM. 1991. Substrate flow in the rhizosphere. The Rhizosphere and Plant Growth.

40. Korenblum E, Dong Y, Szymanski J, Panda S, Jozwiak A, Massalha H, Meir S, Rogachev I, Aharoni A. 2020. Rhizosphere microbiome mediates systemic root metabolite exudation by root-to-root signaling. Proc Natl Acad Sci U S A 117:3874–3883.

41. Han S, Micallef SA. 2016. Environmental metabolomics of the tomato plant surface provides insights on *Salmonella enterica* colonization. Appl Environ Microbiol 82:3131–3142.

42. Gu Y, Wei Z, Wang X, Friman V-P, Huang J, Wang X, Mei X, Xu Y, Shen Q, Jousset A. 2016. Pathogen invasion indirectly changes the composition of soil microbiome via shifts in root exudation profile. Biol Fertil Soils 52:997–1005.

43. Canarini A, Kaiser C, Merchant A, Richter A, Wanek W. 2019. Root exudation of primary metabolites: Mechanisms and their roles in plant responses to environmental stimuli. Front Plant Sci 10:157.

44. Williams A, de Vries FT. 2020. Plant root exudation under drought: implications for ecosystem functioning. New Phytol 225:1899–1905.

45. Carvalhais LC, Dennis PG, Fan B, Fedoseyenko D, Kierul K, Becker A, von Wiren N, Borriss R. 2013. Linking plant nutritional status to plant-microbe interactions. PLoS One 8:e68555.

46. Zhalnina K, Louie KB, Hao Z, Mansoori N, da Rocha UN, Shi S, Cho H, Karaoz U, Loqué D, Bowen BP, Firestone MK, Northen TR, Brodie EL. 2018. Dynamic root exudate chemistry and microbial substrate preferences drive patterns in rhizosphere microbial community assembly. Nature Microbiology 10.1038/s41564-018-0129-3.

47. Chaparro JM, Badri DV, Vivanco JM. 2014. Rhizosphere microbiome assemblage is affected by plant development. ISME J 8:790–803.

48. Cerna-Vargas JP, Santamaría-Hernando S, Matilla MA, Rodríguez-Herva JJ, Daddaoua A, Rodríguez-Palenzuela P, Krell T, López-Solanilla E. 2019. Chemoperception of Specific Amino Acids Controls Phytopathogenicity in *Pseudomonas syringae* pv. tomato. MBio 10.

49. Antúnez-Lamas M, Cabrera-Ordóñez E, López-Solanilla E, Raposo R, Trelles-Salazar O, Rodríguez-Moreno A, Rodríguez-Palenzuela P. 2009. Role of motility and chemotaxis in the pathogenesis of *Dickeya dadantii* 3937 (ex Erwinia chrysanthemi 3937). Microbiology 155:434–442.

50. Webb BA, Compton KK, Del Campo JSM, Taylor D, Sobrado P, Scharf BE. 2017. *Sinorhizobium meliloti* chemotaxis to multiple amino acids is mediated by the chemoreceptor McpU. Mol Plant Microbe Interact 30:770–777.

51. Barbour WM, Hattermann DR, Stacey G. 1991. Chemotaxis of *Bradyrhizobium japonicum* to soybean exudates. Appl Environ Microbiol 57:2635–2639.

52. Gupta Sood S. 2003. Chemotactic response of plant-growth-promoting bacteria towards roots of vesicular-arbuscular mycorrhizal tomato plants. FEMS Microbiol Ecol 45:219–227.

53. Zhou B, Szymanski CM, Baylink A. 2023. Bacterial chemotaxis in human diseases. Trends Microbiol 31:453–467.

54. Gavira JA, Gumerov VM, Rico-Jimenez M, Petukh M, Upadhyay AA, Ortega A, Matilla MA, Zhulin Igor B, Krell T. 2020. How Bacterial Chemoreceptors Evolve Novel Ligand Specificities. MBio 11:1–15.

55. Nishiyama S-I, Suzuki D, Itoh Y, Suzuki K, Tajima H, Hyakutake A, Homma M, Butler-Wu SM, Camilli A, Kawagishi I. 2012. Mlp24 (McpX) of *Vibrio cholerae* implicated in pathogenicity functions as a chemoreceptor for multiple amino acids. Infect Immun 80:3170–3178.

56. Yang Y, M Pollard A, Höfler C, Poschet G, Wirtz M, Hell R, Sourjik V. 2015. Relation between chemotaxis and consumption of amino acids in bacteria: Amino acid chemotaxis and consumption. Mol Microbiol 96:1272–1282.

57. Lane MC, Lloyd AL, Markyvech TA, Hagan EC, Mobley HLT. 2006. Uropathogenic *Escherichia coli* strains generally lack functional Trg and Tap chemoreceptors found in the majority of *E. coli* strains strictly residing in the gut. J Bacteriol 188:5618–5625.

58. Remenant B, de Cambiaire J-C, Cellier G, Jacobs JM, Mangenot S, Barbe V, Lajus A, Vallenet D, Medigue C, Fegan M, Allen C, Prior P. 2011. *Ralstonia syzygii*, the Blood Disease Bacterium and some Asian *R. solanacearum* strains form a single genomic species despite divergent lifestyles. PLoS One 6:e24356.

59. Hasegawa T, Kato Y, Okabe A, Itoi C, Ooshiro A, Kawaide H, Natsume M. 2019. Effect of Secondary Metabolites of Tomato (*Solanum lycopersicum*) on Chemotaxis of *Ralstonia solanacearum*, Pathogen of Bacterial Wilt Disease. J Agric Food Chem 67:1807–1813.

60. Hida A, Oku S, Nakashimada Y, Tajima T, Kato J. 2017. Identification of boric acid as a novel chemoattractant and elucidation of its chemoreceptor in *Ralstonia pseudosolanacearum* Ps29. Sci Rep 7:8609.

61. Sabbagh CRR, Carrere S, Lonjon F, Vailleau F, Macho AP, Genin S, Peeters N. 2019. Pangenomic type III effector database of the plant pathogenic *Ralstonia* spp. PeerJ 7:e7346.

62. Aoun N, Georgoulis SJ, Avalos JK, Grulla KJ, Miqueo K, Tom C, Lowe-Power TM. 2024. A pangenomic atlas reveals eco-evolutionary dynamics that shape type VI secretion systems in plant-pathogenic *Ralstonia*. MBio 15:e0032324.

63. Lefeuvre P, Cellier G, Remenant B, Chiroleu F, Prior P. 2013. Constraints on genome dynamics revealed from gene distribution among the *Ralstonia solanacearum* species. PLoS One 8:e63155.

64. Mattingly HH, Kamino K, Ong J, Kottou R, Emonet T, Machta BB. 2024. *E. coli* do not count single molecules. bioRxiv. 10.1101/2024.07.09.602750.

65. Sneddon MW, Pontius W, Emonet T. 2012. Stochastic coordination of multiple actuators reduces latency and improves chemotactic response in bacteria. Proc Natl Acad Sci U S A 109:805–810.

66. Fernandes AS, Campos KF, de Assis JCS, Gonçalves OS, Queiroz MV de, Bazzolli DMS, Santana MF. 2024. Investigating the impact of insertion sequences and transposons in the genomes of the most significant phytopathogenic bacteria. Microb Genom 10.

67. Stanier RY, Palleroni NJ, Doudoroff M. 1966. The aerobic pseudomonads: a taxonomic study. J Gen Microbiol 43:159–271.

68. Chen I-MA, Chu K, Palaniappan K, Pillay M, Ratner A, Huang J, Huntemann M, Varghese N, White JR, Seshadri R, Smirnova T, Kirton E, Jungbluth SP, Woyke T, Eloe-Fadrosh EA, Ivanova NN, Kyrpides NC. 2019. IMG/M v.5.0: an integrated data management and comparative analysis system for microbial genomes and microbiomes. Nucleic Acids Res 47:D666–D677.

69. Paysan-Lafosse T, Blum M, Chuguransky S, Grego T, Pinto BL, Salazar GA, Bileschi ML, Bork P, Bridge A, Colwell L, Gough J, Haft DH, Letunić I, Marchler-Bauer A, Mi H, Natale DA, Orengo CA, Pandurangan AP, Rivoire C, Sigrist CJA, Sillitoe I, Thanki N, Thomas PD, Tosatto SCE, Wu CH, Bateman A. 2023. InterPro in 2022. Nucleic Acids Res 51:D418–D427.

70. Parks DH, Imelfort M, Skennerton CT, Hugenholtz P, Tyson GW. 2015. CheckM: assessing the quality of microbial genomes recovered from isolates, single cells, and metagenomes. Genome Res 25:1043–1055.

71. Arkin AP, Cottingham RW, Henry CS, Harris NL, Stevens RL, Maslov S, Dehal P, Ware D, Perez F, Canon S, Sneddon MW, Henderson ML, Riehl WJ, Murphy-Olson D, Chan SY, Kamimura RT, Kumari S, Drake MM, Brettin TS, Glass EM, Chivian D, Gunter D, Weston DJ, Allen BH, Baumohl J, Best AA, Bowen B, Brenner SE, Bun CC, Chandonia J-M, Chia J-M, Colasanti R, Conrad N, Davis JJ, Davison BH, DeJongh M, Devoid S, Dietrich E, Dubchak I, Edirisinghe JN, Fang G, Faria JP, Frybarger PM, Gerlach W, Gerstein M, Greiner A, Gurtowski J, Haun HL, He F, Jain R, Joachimiak MP, Keegan KP, Kondo S, Kumar V, Land ML, Meyer F, Mills M, Novichkov PS, Oh T, Olsen GJ, Olson R, Parrello B, Pasternak S, Pearson E, Poon SS, Price GA, Ramakrishnan S, Ranjan P, Ronald PC, Schatz MC, Seaver SMD, Shukla M, Sutormin RA, Syed MH, Thomason J, Tintle NL, Wang D, Xia F, Yoo H, Yoo S, Yu D. 2018. KBase: The United States department of energy systems biology knowledgebase. Nat Biotechnol 36:566–569.

72. Letunic I, Bork P. 2024. Interactive Tree of Life (iTOL) v6: recent updates to the phylogenetic tree display and annotation tool. Nucleic Acids Res 52:W78–W82.

73. Gilchrist CLM, Chooi Y-H. 2021. Clinker & clustermap.Js: Automatic generation of gene cluster comparison figures. Bioinformatics 37:2473–2475.

74. Seemann T. 2014. Prokka: rapid prokaryotic genome annotation. Bioinformatics 30:2068–2069.

75. Krzywinski M, Schein J, Birol I, Connors J, Gascoyne R, Horsman D, Jones SJ, Marra MA. 2009. Circos: an information aesthetic for comparative genomics. Genome Res 19:1639–1645.

76. Wolfe AJ, Berg HC. 1989. Migration of bacteria in semisolid agar. Proc Natl Acad Sci U S A 86:6973–6977.

77. Miller JH. 1975. Miller, J.H. (1975) Experiments in molecular genetics. Cold Spring Harbor Laboratory, 359 Cold Spring Harbor, N.Y. Cold Spring Harbor Laboratory 359.

78. Salanoubat M, Genin S, Artiguenave F, Gouzy J, Mangenot S, Arlat M, Billault A, Brottier P, Camus JC, Cattolico L, Chandler M, Choisne N, Claudel-Renard C, Cunnac S, Demange N, Gaspin C, Lavie M, Moisan A, Robert C, Saurin W, Schiex T, Siguier P, Thébault P, Whalen M, Wincker P, Levy M, Weissenbach J, Boucher CA. 2002. Genome sequence of the plant pathogen *Ralstonia solanacearum*. Nature 415:497–502.

79. Ailloud F, Lowe TM, Robène I, Cruveiller S, Allen C, Prior P. 2016. In planta comparative transcriptomics of host-adapted strains of *Ralstonia solanacearum*. PeerJ 4:e1549.

80. Ailloud F, Lowe T, Cellier G, Roche D, Allen C, Prior P. 2015. Comparative genomic analysis of *Ralstonia solanacearum* reveals candidate genes for host specificity. BMC Genomics 16:1–11.

81. Castañeda A, Reddy JD, El-Yacoubi B, Gabriel DW. 2005. Mutagenesis of all eight avr genes in *Xanthomonas campestris* pv. campestris had no detected effect on pathogenicity, but one avr gene affected race specificity. Mol Plant Microbe Interact 18:1306–1317.

